# Geographical variation in mitogenomes of the largetooth sawfish *Pristis pristis*: challenges and perspectives for conservation efforts

**DOI:** 10.1101/2025.02.22.639618

**Authors:** Alan Érik S. Rodrigues, Rafaela Maria S. Brito, Patricia Charvet, Vicente V. Faria, Mariano Cabanillas-Torpoco, Alexandre Aleixo, Tibério César T. Burlamaqui, Luis Fernando da S. Rodrigues-Filho, Angelico Asenjo, Raquel Siccha-Ramirez, Jorge Luiz S. Nunes, Hugo J. de Boer, José Cerca, Quentin Mauvisseau, Jonathan S. Ready, João Bráullio L. Sales

## Abstract

Sawfishes (Pristidae) have been severely impacted by coastal development and unregulated fisheries and are considered Critically Endangered by the IUCN Red List. Environmental DNA (eDNA) analyses have shown potential for monitoring elasmobranch species, with various studies focusing on using species-specific approaches to detect *Pristis* species. However positive detection using existing probes has not been confirmed in some geographic regions where they would be expected. Here, we aimed to test whether mutations at key sites have been detrimental to species-specific detection of *P. pristis* (Linnaeus, 1758) using the existing probe set. To test this hypothesis mitogenomes were assembled that were found to follow the typical pattern of vertebrate mitogenomic organization. Phylogenetic trees showed similar topologies and confirmed geographic mitochondrial variation in *P. pristis*. Mismatches for the published 12S species-specific probe set for *P. pristis* were identified that prevent amplification of positive control samples from Brazil. However, ddPCR detection of the positive control was possible using a newly designed species-specific probe set. This study highlights how geographical variation can severely impact the success of generally applying species-specific detection systems developed based on data from only one geographical region. The new mitogenomes and species-specific probe set developed here may also contribute to improving the potential to map and monitor these Critically Endangered species across the globe.

**Graphical abstract.**
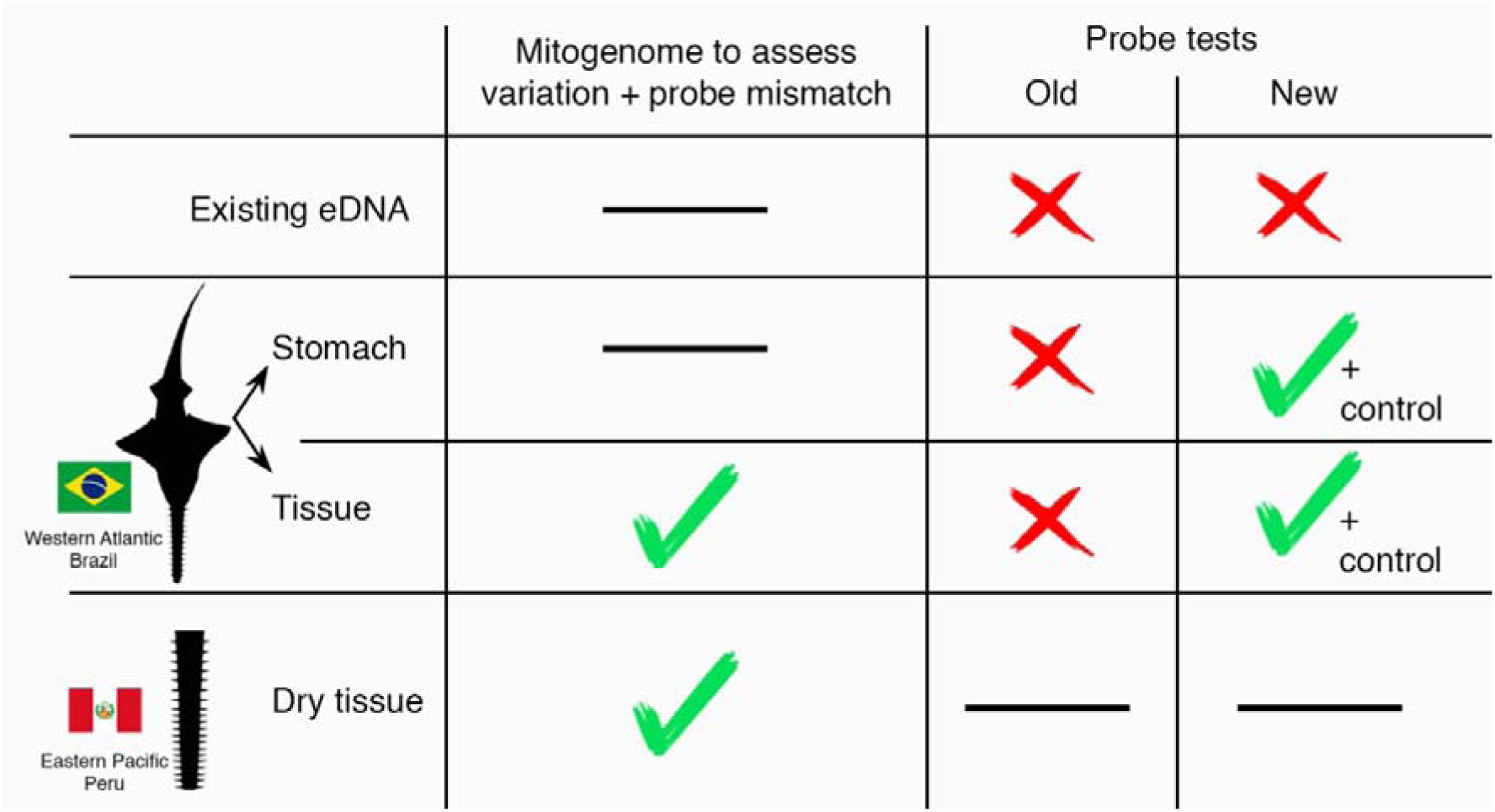

## 1. Introduction

Sawfishes (Pristidae) were once common in soft-bottom habitats of shallow, warm waters worldwide, often holding significant cultural importance as reflected in mythology and folklore (Robillard and Séret, 2006; Harrison and Dulvy, 2014; McDavitt, 2014; Moore, 2017; Cabanillas-Torpoco et al., 2023). However, over the past three decades, their populations have been severely impacted by coastal development and unregulated fisheries (Cavanagh et al., 2005; Dulvy et al., 2014). Today, sawfishes have largely disappeared from vast areas at local, regional, and global scales (Fernandez□Carvalho et al., 2014; Moore, 2015; Dulvy et al., 2016; Leeney and Downing, 2016). They are now considered the most threatened family of elasmobranchs globally (Dulvy et al., 2014), comprising five extant species: *Anoxypristis cuspidata* (Latham, 1794), *Pristis pristis* (Linnaeus, 1758), *Pristis pectinata* Latham, 1794, *Pristis zijsron* Bleeker, 1851, and *Pristis clavata* Garman, 1906 (Faria et al., 2013). All are currently listed as “Critically Endangered” on the IUCN Red List (Carlson et al., 2022; Espinoza et al., 2022; Grant et al., 2022; Harry et al., 2022; Haque et al., 2023).

The application of molecular methods in recent years has significantly advanced the study of elasmobranchs, leading to the description of new species (Naylor et al. 2012, White et al., 2013), the reorganization and resurrection of families, and the reassignment of taxa (Last et al., 2016; White and Naylor, 2016; Rodrigues-Filho et al., 2023). These methods have also revealed the presence of cryptic species (Sales et al., 2019; Gales et al., 2024; Petean et al., 2024). Among sawfishes, the Common Sawfish (*P. pristis*) is one of the most iconic species and serves as the type species of its genus (Bigelow and Schroeder, 1953). Originally described by Linnaeus (1758) as having a distribution “in Europa”, the species is now considered to have a circumglobal, presence in tropical and warm temperate seas, after a taxonomy review based on molecular and morphological evidence (Faria et al., 2013).

Environmental DNA (eDNA) analyses have shown potential for monitoring elasmobranch species using metabarcoding (Boussarie et al., 2018; West et al., 2020; Budd et al., 2021; de la Hoz Schilling et al., 2024), as well as using species-specific detection methods that focus on *Pristis* species (Simpfendorfer et al., 2016; Bonfil et al., 2021; Cooper et al., 2021). However, species-specific identification through eDNA detection can produce false negatives if local geographic lineages have sequences that vary at the site of the specifically designed primers/probe (Wilcox et al., 2015). The species-specific primers and probe set available for *P. pristis* detection were developed from and for the Australian population (Simpfendorfer et al., 2016), and because there are indications of geographical population structure within *P. pristis* (Faria et al., 2013; Feutry et al., 2015), it is important to consider whether the lack of detection of this species in other regions of the world (Bonfil et al., 2021, Rodrigues et al., unpublished results) could be due to mismatches between the primers/probe and the mitogenomic sequences in the local populations to which they should anneal.

Here, we aimed to test whether mutations at key sites have been detrimental to species-specific detection of *P. pristis* in the Eastern Pacific and Western Atlantic using the existing probe set. To test this hypothesis, mitogenomes were successfully assembled as a form of superbarcode (Crampton-Platt et al., 2016) to provide regional reference sequences for *P. pristis*, potentially revealing the need to develop and test alternative regional species-specific primers for *P. pristis*.

## 2. Methods

### 2.1. Sampling

A sample of muscle tissue and stomach contents of a juvenile individual *P. pristis* from the Western Atlantic (specifically from 3°23’05’ S 44° 48’ 40’ W, Bonfim do Arari, upper part of the estuary of São Marcos Bay, Maranhão, Brazil), that was caught in a fishing net and donated by fishermen to JLSN. The sample collection and transport permit (“Sistema de Autorização e Informação em Biodiversidade” - SISBIO) for the present study was license 60306-4, and the genetic patrimony (“Sistema Nacional de Gestão ao Patrimônio Genético e do Conhecimento Tradicional Associado” - SisGen) license was A9851C4. A small sample of cartilage was also taken from an old dry *P. pristis* rostrum from Talara, Eastern Pacific in Peru, a historical (1960, 65 years ago) personal item belonging to the family of MC-T, which was fished and commercialized before the species was granted legal protected status.

### 2.2. DNA extraction and mitogenomic sequencing

The tissue samples were processed in independent laboratories in Brazil and Peru. All procedures were performed following initial decontamination of all materials and surfaces using bleach and UV light exposure. Tissue and cartilage samples were extracted using an adapted (addition of 20 µl proteinase K for tissue lysis) CTAB protocol (Doyle and Doyle, 1987) in decontaminated laminar flow cabinets. DNA was quantified in Qubit 4 Fluorometer (Thermo Fisher Scientific Inc., MA, USA). Libraries were prepared following standard Illumina protocols using genomic DNA concentration of 20 ng and tagmentation times varying from 20 to 25 minutes, followed by fragment analysis using a Tapestation 4200 (Agilent, CA, USA). The libraries were sequenced on Illumina NextSeq 2000 platforms (Illumina, San Diego, CA, USA) using NextSeq 1000/2000 P2 reagents (100 cycles) to generate 300 bp paired-end reads (insert size varying from 459 to 526 bp) resulting in ∼6x genome coverage.

To obtain a positive mixed-species control sample (trace DNA of *P. pristis* along with DNA of other species), the stomach contents were separated from the ethanol preservative by three cycles of washing and centrifugation with UV sterilized ultrapure water. Four 650 μl subsample replicates and one negative control (non-template ultrapure water) were prepared and stored in separate microcentrifuge tubes (Rosa et al., 2024).

### 2.3. Mitogenome assembly and annotation

Quality of the generated reads was checked using FastQC v.0.12.1 (Andrews, 2010), then trimmed to remove adapters and exclude reads shorter than 70 base pairs. Complete mitogenomes were assembled with NovoPlasty v.4.3.5 (Dierckxsens, et al., 2016) and annotated using the MITOS2 web server v.2.1.7 (Donath et al., 2019). All tRNAs structures were identified and visualized with tRNAscan (Chan and Lowe, 2019) with default parameters. To determine accurate limits of all ribosomal units, protein-coding genes and secondary structure we ran an additional annotation with MITOFish v.4.01 (Zhu et al., 2023), then aligned and visually checked against an existing mitogenome of *P. pristis* from Australia (NC_039438.1 on GenBank) using Geneious Prime bioinformatic software v. 2024.0.5 (https://www.geneious.com).

Mitogenomes were drawn in a full circle using CGviewer Server (Grant et al., 2023). The GC skew was established and added to the graph using the following calculation (G − C)/(G + C). The relative synonymous codon usage (RSCU) values were calculated in MEGA X (Kumar et al., 2021) and the nucleotide diversity based on 13 PCGs of *Pristis* genus with 100 bp sliding window and each step of 25 bp, was estimated with DNAsp6 (Rozas et al., 2017), the results of both analyses were plotted with RStudio (R Core Team, 2023) using the ggplot2 package (Wickham, 2006).

### 2.4. Phylogenomic analyses

We built a dataset containing all complete mitogenomes available on GenBank for Rhinopristiformes and included complete mitogenomes for a holocephalan (*Callorhinchus milii* Bory de Saint-Vincent, 1823) and an actinopterygian (*Tor putitora* (Hamilton, 1822)) (Table 1) as the same outgroups used by Wang et al. (2023). Only the 12 H-strand PCGs and the rRNA genes were used. Although the ND6 is a coding gene, it is on the L-Strand where there is a distinct asymmetry of base composition (Miya and Nishida, 2015), and it was therefore not included in the analysis. The sequences were aligned with standard MUSCLE alignment algorithm (Edgar, 2004) and visually revised on Geneious Prime bioinformatic software. To improve accuracy of phylogenetic inference, we excluded the start and stop codons of all H-strand PCGs due to their highly conserved structure, as well as the third codon position due to the high variation that can give rise to synonymous mutations that do not have functional impact (Näsvall et al., 2023). In addition, the alignment of the H-strand PCGs was split in the first and second codon position. We used Gblock v.0.91b (Castresana, 2000) to exclude ambiguous alignments for ribosomal subunits. The segments of rRNA units and H-strand PCGs were concatenated with Phylosuite v.1.2.3 (Xiang et al., 2023). After that we defined the rRNA units, first and second codons as partitions and tested whether these partitions should be analyzed using distinct evolutionary models using Partition Finder v.2.1.1 (Lanfear et al., 2017). We used ModelFinder (Kalyaanamoorthy et al., 2017) to determine the best edge-equal evolutionary model for each partition as: rRNAs (TIM2+F+R3), first codon (GTR+F+I+G4) and second codon position (TIM3+F+I+R2). A Maximum Likelihood tree was generated using 1.000.000 pseudoreplicates and the Ultrafast Bootstrap algorithm (Minh et al., 2013) on IQ-tree v.2.3.4 (Minh et al., 2020) with an *abayes* approximation test (Anisimova et al., 2011). Furthermore, a Bayesian Inference analysis based on the same dataset and partition model was conducted with 2 parallel runs with 1.000.000 generations, in which 25% of the initial tree was discarded as burn-in. Both trees generated were visualized and annotated using Figtree software v.1.4.4 (Rambaut, 2018).

**Table1.** List of mitogenomes used for phylogenetic analyses. The codes of the mitogenomes from this study are represented in bold.

| Family | Specie | Length(bp) | GenBank code |
| --- | --- | --- | --- |
| Callorhinchidae | *Callorhinchus milii* | 16,769 | NC_014285 |
| Cyprinidae | *Tor putitora* | 16,576 | NC_021755 |
| Glaucostegidae | *Glaucostegus granulatus* | 16,547 | MN783017 |
| Pristidae | *Anoxypristis cuspidata* | 17,243 | NC_026307 |
| Pristidae | *Pristis clavata* | 16,804 | NC_022821 |
| Pristidae | *Pristis pectinata* | 16,802 | NC_027182 |
| Pristidae | *Pristis pristis* | 16,912 | NC_039438 |
| Pristidae | *Pristis pristis* (Western Atlantic) | 16,807 | **PV053514** |
| Pristidae | *Pristis pristis* (Eastern Pacific) | 16,914 | **PV053515** |
| Pristidae | *Pristis zijsron* | 16,804 | MH005927 |
| Rhinidae | *Rhina ancylostoma* | 17,217 | NC_030215 |
| Rhinidae | *Rhynchobatus australiae* | 16,804 | NC_030254 |
| Rhinidae | *Rhynchobatus djiddensis* | 16,799 | NC_066688 |
| Rhinidae | *Rhynchobatus laevis* | 16,560 | NC_047241 |
| Rhinobatidae | *Acroteriobatus annulatus* | 16,773 | NC_068897 |
| Rhinobatidae | *Acroteriobatus blochii* | 16,771 | NC_068898 |
| Rhinobatidae | *Rhinobatos hynnicephalus* | 16,776 | NC_022841 |
| Rhinobatidae | *Rhinobatos schlegelii* | 16,780 | NC_023951 |
| Trygonorrhinidae | *Zapteryx exasperata* | 17,310 | NC_024937 |

### 2.5. Divergence time analyses

The divergence time to The Most Recent Common Ancestor (TMRCA) was estimated based on a Bayesian MCC tree that was produced in BEAST 2 (Bouckaert et al., 2019) using the dataset above. The Yule speciation prior was used for the tree prior, modeled with an uncorrelated lognormal relaxed clock (Drummond et al., 2006) with GTR+G model. We followed the same calibration priors of Wang et al. (2023): A-159.2 million years ago (Mya) for the common ancestor of Rhinopristiformes; B-85.6 Mya for the common ancestor of the genera *Rhina* and *Pristis*; and C-55.3 Mya for the common ancestor of *P. clavata* and *P. pristis.* The MCMC method was used to infer the divergence times with four independent runs with 120 million generations through four simultaneous runs containing four chains (one cold and three heated) with sampling performed every 1000 generations. Only runs with ESS values equal to or greater than 200 for all marginal parameters were used. The log-likelihood files generated from each run were viewed in Tracer v. 1.5 (Rambaut and Drummond, 2009), to check if the ESS values were equal to, or greater than, 200 after discarding 10% of the trees as burn-in. The consensus MCC tree was then generated using the TreeAnnotator v. 1.4 (Drummond and Rambaut, 2007).

### 2.6. Species-specific eDNA probes

The previously published species-specific primers and probes targeting *P. pristis* (Cooper et al., 2021; Cooper et al., 2022) were aligned with mitogenomes available in GenBank and others generated in this study using Geneious Pro R10 (Kearse et al., 2012) with the “multiple alignment” function to identify potential mismatches susceptible to lead to false negative detection. Additionally, sequences from the target species as well as from closely related Pristidae species and potentially co-occurring (in the western Atlantic) and/or closely related elasmobranch species deposited in GenBank (Appendix A, Table A.1) were downloaded and also aligned using the “multiple alignment” function available on Geneious Pro R10 (Kearse et al., 2012). Conserved fragments within the *P. pristis* sequences, showing variation with non-target species, were identified, and a new set of species-specific primers and probe targeting a 151 bp fragment of the mitochondrial 12S region of *P. pristis* were designed using the “primers” design function of Geneious (Brys et al., 2020; Mauvisseau et al., 2021; Bommerlund et al., 2023). In addition to the visual alignment with sequences from target and non-target species used to design the assay, its specificity was further confirmed using the NCBI primer-blast function, with both “Forward and Reverse primers”, “Probe and Reverse primer”, and “Forward primer and reverse Probe fragment” combinations. *In-vitro* testing was conducted using trace DNA extracted from stomach contents of the juvenile sample from the western Atlantic as well as previously collected predominantly near-shore eDNA samples from the Atlantic coast of Northern Brazil (Appendix A, Table A.2). This was both to confirm the amplification of target material in mixed-species DNA samples (traces of *Pristis* DNA within its stomach contents) using the newly developed assay, and to assess as best as possible the assay’s specificity. It was expected that a significant proportion of the many pre-existing near-shore eDNA sampling locations should not show presence of *P. pristis* and therefore act as negative controls by registering the absence of amplification of local non-target species diversity. This was done to mitigate challenges in obtaining genetic material needed for further laboratory validation because all Pristidae are CITES listed, which limits sharing genetic material across countries.

Samples were analyzed using both the existing species-specific primer and probe set (Cooper et al., 2021) as well as the newly developed primer and probe set on a Bio-Rad QX200 ddPCR System. ddPCR reactions were performed in a 20 µl final volume, consisting of 10 µl Bio-Rad ddPCR supermix for probes (no dUTP), 0.75µM of forward and reverse primers, 0.375 µM of probe, 5.0 µl of ddH2O and 2 µl template DNA. For each reaction, droplets were generated using a DG8 Droplet Generator Cartridge and 70 µl of Droplet Generation Oil for Probes on a QX200 Droplet Generator (Bio-Rad), and a final 40 µl volume of droplets for each reaction was transferred to a ddPCR 96-well plate. End-point PCR amplifications were performed on a BioRad CFX96 Real-Time System (Bio-Rad Laboratories, California, United States). PCR conditions were as follows: 10 min at 95°C, followed by 40 cycles of denaturation for 30 s at 94°C and annealing at 55°C for 1 min, with ramp rate of 2°C/s, followed by 10 min at 98°C and a hold at 8°C for our newly developed primers/probe set. Similar PCR conditions with an annealing temperature of 60°C were used with the primers/probe set described in Cooper et al. (2021). Droplets were read on a QX200 droplet reader (Bio-Rad), and quantification was achieved using the Bio-Rad QuantaSoft software (v.1.7.4.0917). Thresholds for positive signals were determined according to QuantaSoft software instructions, and all droplets above the fluorescence threshold were counted as positive events, those below it being counted as negative events.

## 3. Results

### 3.1. Mitogenomic variation

The new complete mitogenome of *P. pristis* from the Western Atlantic has total length of 16,807 bp, with the following nucleotide composition: T(U), 28.3%; C, 26.6%; A, 32.1%; G, 13.0%, resulting in an A+T content of 60% and C+G of 40%. In comparison, the mitogenome from *P. pristis* from the Eastern Pacific is 16,914 bp long with the following composition: T(U), 28.2%; C, 26.7%; A, 32.1%; G, 13.1%, also showing an A+T and C+G proportion of 60% and 40%, respectively (Fig. 1). The mitochondrial arrangement also follows the typical vertebrate pattern, containing 2 rRNA genes, 13 protein-coding genes, 22 tRNA genes (see tRNAs structures in Appendix B), and a non-coding control region (*D-loop*). However, individuals from the Eastern Pacific and Australia display a distinct feature: both have 23 tRNA genes due to the duplication of the tRNA-Pro located between the tRNA-Thr and the control region. In contrast, the mitogenome of *P. pristis* from the Western Atlantic lacks this duplication, marking a significant distinction compared to the other *P. pristis* mitogenomes described so far (Fig. 1). RSCU values showed little variation following sawfish patterns and nucleotide diversity indicates somewhat less variation in the three COX gene units compared to ATPase and NADH units at the genus level (see Appendix B, Fig. B.1-B.3).

**Fig. 1.**
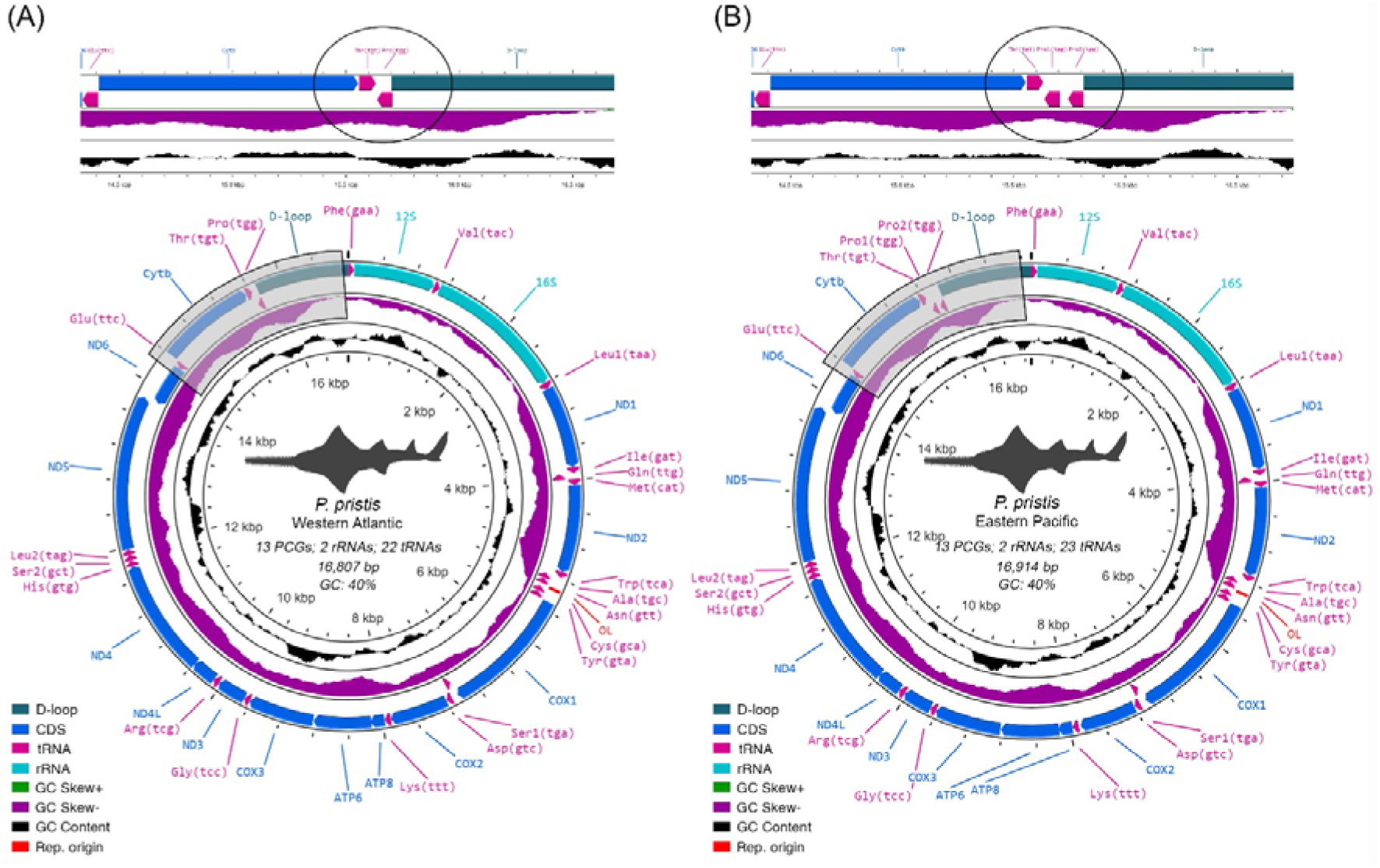
Mitogenomes of new and geographically distinct samples of *P. pristis* with highlighted structural differences: (A) Mitogenome from the Western Atlantic sample. (B) Mitogenome from the Eastern Pacific sample.

### 3.2. Phylogenetic relationships of Rhinopristiformes and divergence time estimations

All branches of the phylogenetic trees inferred from the 12 H-strand PCGs and ribosomal units of all Rhinopristiform species were well supported (Appendix C, Fig. C.4A, 4B), confirming the monophyly of the five families analyzed, diverging only in the position of *Glaucostegus granulatus* (Cuvier, 1829). However, the topology obtained reinforces the complete resolution of the family Pristidae. Moreover, within *P. pristis*, it confirms evidence of geographic variation between the Australian, Eastern Pacific, and Western Atlantic samples supported by both ML and Bayesian analyses (Appendix C, Fig. C.4A, 4B).

The Maximum Clade Consensus tree (Fig. 2) shows a similar topology to that of Wang et al. (2023). However, we recovered differences in the estimated time since divergence between the outgroups utilized, estimated as ∼150.85 mya (HPD 95%: 155.20- 182.41 mya, Fig. 2) and the most common recent ancestor of outgroups and Rhinopristiformes which was estimated at ∼168.13 mya (HPD 95%: 155.20-182.441 mya) instead of ∼310.93 and ∼361.83 mya, respectively, predicted by Wang et al. (2023). The common ancestor of all Rhinopristiformes in our simulation was inferred to exist ∼114.81 mya (HPD 95%: 103.95-125.24 mya, Fig. 2). The common ancestor of Pristidae (*Anoxypristis* and *Pristis*) was estimated to exist ∼71,74 mya (HPD 95%: 66.06-77.21 mya, Fig. 2), compatible with the estimate of Wang et al, (2023) (∼76.42 mya) considering the confidence intervals. The divergence estimates for *P. pristis* samples from the Western Atlantic, Eastern Pacific and Australian regions reveal that the variation in mitogenomic sequences represents a historical process.

**Fig. 2.**
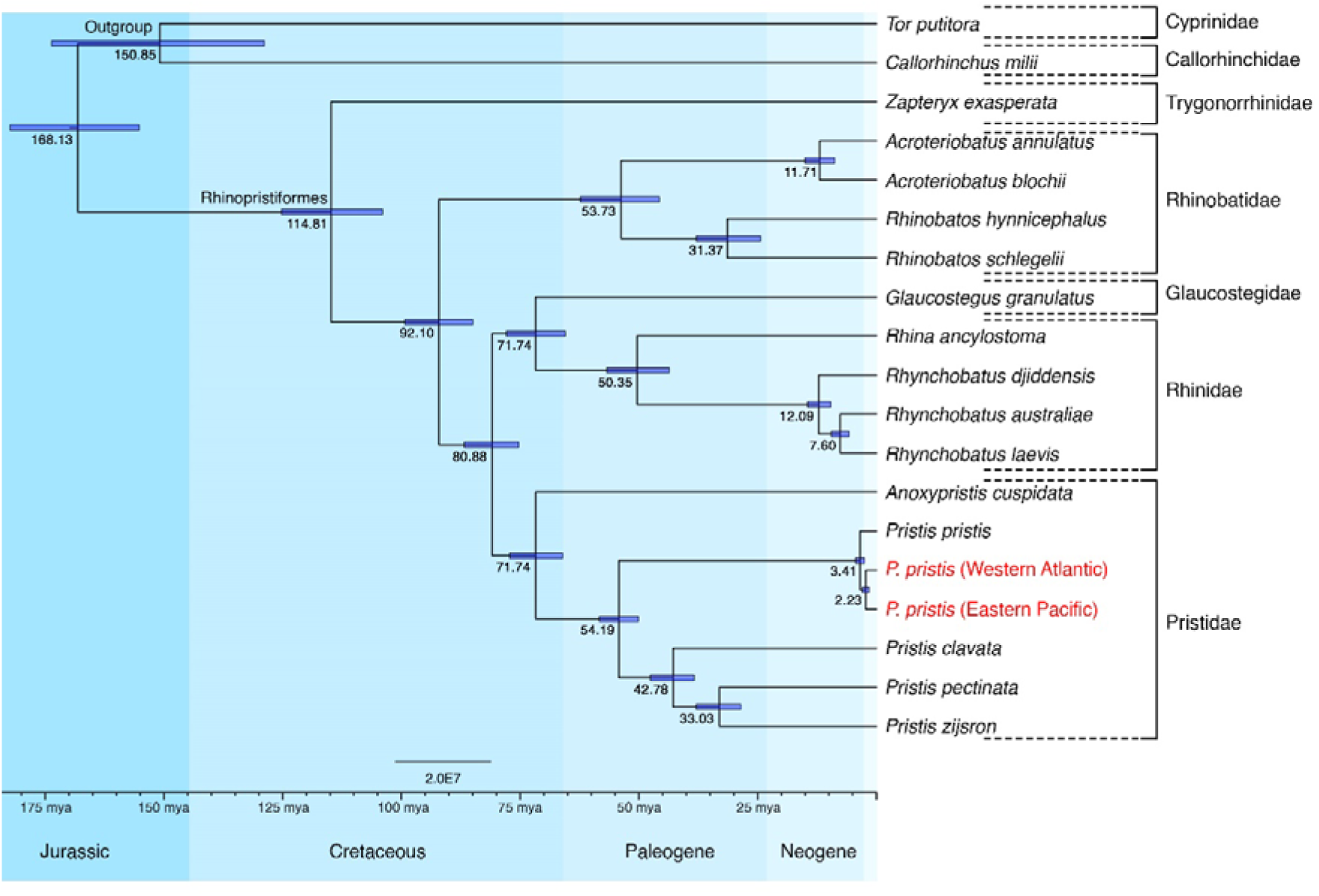
Maximum clade consensus tree including divergence time estimates for Rhinopristiformes generated using the 12 H-strand PCGs and rRNA genes inferred using BEAST2. The new P. pristis from this study are in red.

### 3.3. Species-specific eDNA probes

The alignment of the previously published 12S *P. pristis* species-specific primers and probe set (Cooper et al., 2021) with the new mitogenomes identified one mismatch on the probe sequence for both new mitogenomes (Fig. 3A; Appendix D, Fig. D.2A), one mismatch with the reverse fragment for the mitogenome obtained from the Western Atlantic specimen (Appendix D, Fig. D.2B), and two mismatches for the mitogenome obtained from the Eastern Pacific specimen (Fig. 3B; Appendix D, Fig. D.2B). We were not able to amplify the DNA extracted from the stomach content of the Brazilian specimen using these published primers and probe from Cooper et al. (2021) and obtain a positive ddPCR signal, confirming that the mismatches identified on the probe and reverse primers would lead to false negative results with *P. pristis* occurring in Brazilian waters. However, we were able to amplify the DNA extracted from the stomach content of the Brazilian specimen using the newly designed set of species-specific primers and probe targeting *P. pristis* (Appendix D, Fig. D.6), Forward primer 5’- CCTAAGAAAAAACGAACAGTA -3’, Probe 5’-CACTATTCTGAAACTGGCTC -3’, Reverse primer 5’- GTTTATGTAAGGGGAATATTAT -3’) (Fig. D.3-D.5; Appendix D). Furthermore, the analysis of eDNA samples collected in the biodiversity rich marine systems of the Northern Brazilian coastline led to no detectable PCR amplification using the ddPCR platform and newly developed assay, confirming that this assay did not amplify non-target and/or co-occurring species in this region which could lead to false positive results.

**Fig. 3.**
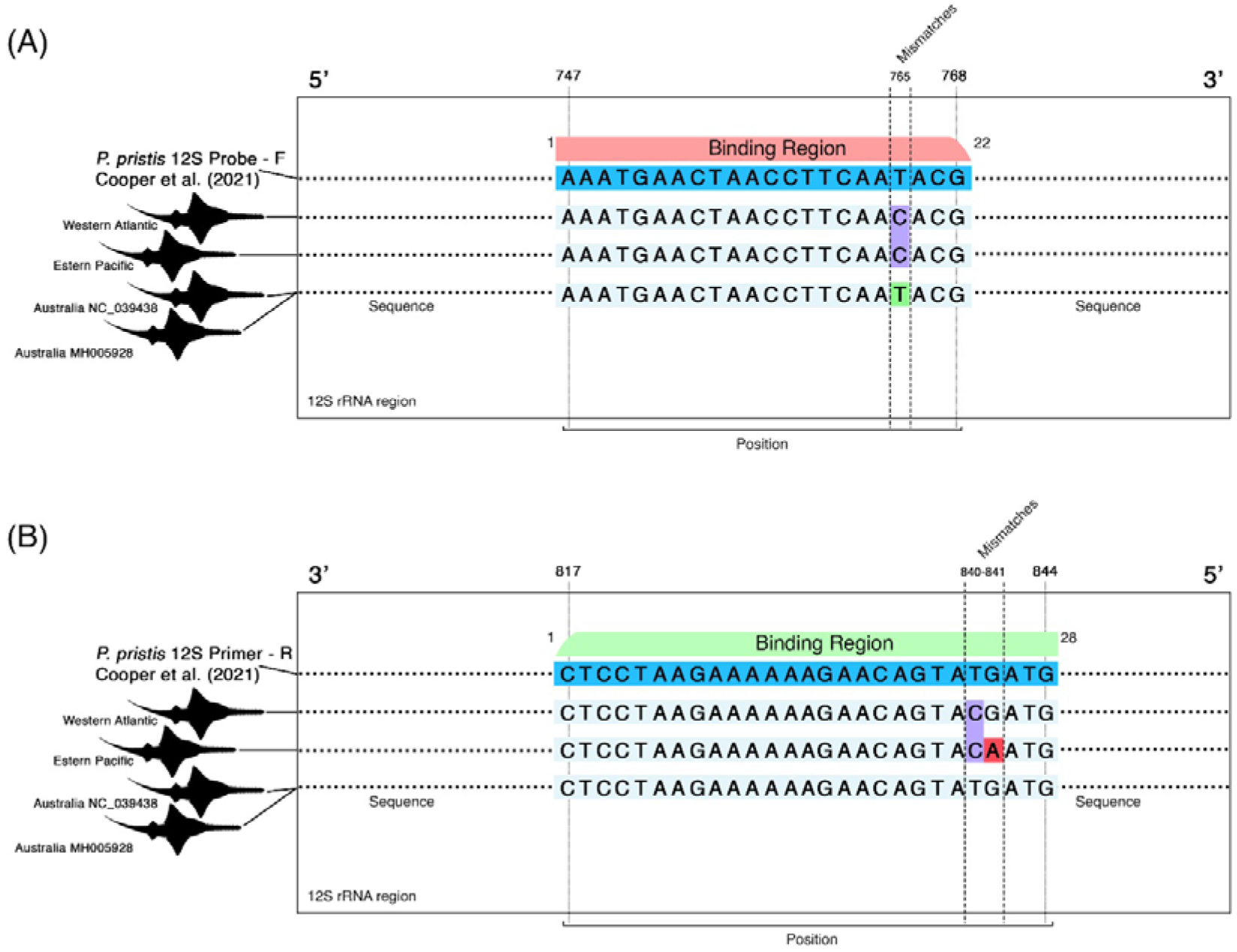
Local alignment of the existing (Cooper et al., 2021) 12S species-specific *P. pristis* probe (A) and reverse primer (B) with *P. pristis* mitogenomes including the two new mitogenomes produced in this study for the Western Atlantic and Eastern Pacific samples as well as two Australian mitogenomes available in GenBank showing the mismatch of nucleotides for both the Western Atlantic and Eastern Pacific mitogenome lineages that prevent primer binding.

## 4. Discussion

### 4.1. Mitogenome comparisons

Both mitogenomes are highly similar to the existing mitochondrial genome of *P. pristis* described by Kyne et al. (2018) with similar GC content, nucleotide composition values and RSCU values to that and to other fish species (Boore, 1999; Kousteni et al., 2021). The slight difference in total mitogenome length between individuals from Australia/Eastern Pacific and Western Atlantic is mainly determined by the deletion of a tRNA-Pro secondary structure duplication in the mitogenome of the sample from the Western Atlantic. However, as there is currently no biological explanation for the loss of this duplication, it is necessary to investigate further, although such rearrangements on the mitochondrial genome have been reported for other bony fish species (Miya and Nishida, 1999), elasmobranchs (Winn et al., 2024; Fee et al., 2025) and other taxa (e.g., amphibians - Zhang et al., 2021, cephalopods - Yokobori et al., 2004; Taite et al., 2023).

### 4.1. Phylogeny and Divergence Time Estimations

The taxonomy of the family Pristidae reised by Faria et al. (2013) based on molecular and morphological data reducing the number of species in the genus *Pristis*, synonymizing *Pristis microdon* and *Pristis perotteti* within *P. pristis* because integrated data did not support the maintenance of these species It was also proposed that *A. cuspidata* was basal to a *Pristis* genus clade, as follows (*A. cuspidata* (*P. pristis* (*P. clavata* (*P. pectinata*, *P. zijsron*)))) (Faria et al., 2013). Kyne et al. (2018) provided the complete mitochondrial genome of *P. pristis* and corroborated the phylogeny established for the family Pristidae (without including *P. zijsron*). Furthermore, Wang et al. (2023) added the complete mitogenome of *P. zijsron*. Our phylogenetic tree was generally consistent with the existing phylogenies using H-strand PCGs from mitogenomes (Kyne et al., 2018; Kousteni et al., 2021; Wang et al., 2023).

*P. pristis* is the sawfish with the broadest geographic distribution. Global geographical structuring of its populations in tropical Western Atlantic, Eastern Atlantic, Eastern Pacific, and Western Indo-Pacific have been detected based on divergence in NADH- 2 mtDNA gene and rostral tooth counts (Faria et al., 2013). Similarly, local geographical structuring has been in northern Australia) have also been detected based on another region of the mtDNA, the D-loop (Phillips et al., 2011) as well as complete mitogenomes (Feutry et al., 2015). Given the previous geographic variation in the mtDNA and the results of this study, additional complete mitogenomes from across the full geographic range will be important to better assess *P. pristis* population structuring. Most of the remaining specimens, particularly isolated rostra, that could provide this coverage are stored dry in museums or private collections, making the process of extracting DNA and DNA sequencing difficult.

### 4.2. Challenges and perspectives for species-specific eDNA monitoring of *P. pristis*

It is clear that the geographical variation described above can severely impact the chances of generally applying species-specific detection systems developed only based on data from one region. The probes previously developed by Cooper et al. (2021) for the Australian population of *P. pristis* did not work for the species-specific detection of *P. pristis* from the Western Atlantic using our positive control (Fig. 4). This confirms that the mismatches in the probe and reverse primer (Fig. 3) associated with these geographic mitogenome variants are most likely responsible for false negatives when attempting to detect these individuals in natural environments of the Western Atlantic when using ddPCR (Bonfil et al., 2021, Rodrigues et al., unpublished results). The new primers and probe developed here successfully amplified the target species’ trace DNA within a mixed-species DNA sample (the stomach content samples from *P. pristis* from Brazil). The lack of positive detection in eDNA samples from the region supports the specificity of the assay and indicates that the search for positive signals of these species will require greater coverage of environmental space and/or improvement of sampling methodologies that increase the chances of recovery of *Pristis* DNA (e.g. stratified sampling of the water column). In support of Lehman et al. (2020), our results further demonstrate that overcoming mitochondrial gene variations among populations of a species (Rubinoff, 2006) requires *in-silico* tests with positive and negative controls to ensure highly reliable validation of probes developed for each local population.

The primers and probe developed in this study may serve as an improvement for eDNA detection approaches and help make them work for local populations and result in more reliable tools for detection and conservation of *P. pristis*, a flagship species in the coastal areas of the Easter Pacific and Western Atlantic. This may be especially important along the Northern coast of Brazil, which is believed to be one of the last population refuges for *P. pristis* in the Western Atlantic (Fernandez-Carvalho et al., 2014; Nunes et al., 2016; Feitosa et al., 2017, Fordham et al., 2018; Fordham et al. 2018). This perspective of an improved ability to detect the critically endangered *P. pristis* in the region (present study) is aligns with Bohmann et al. (2014), who emphasized the crucial importance of mapping species’ occurrence, identifying important ray areas for conservation, and monitoring changes in its distribution over. This approach could also support the implementation of targeted protection measures, enabling more effective conservation efforts of the remaining populations, particularly in areas under anthropogenic pressure, such as fishing and habitat degradation. Moreover, accurate species detection can contribute to global conservation initiatives, aligning with international efforts to prevent its extinction (Dulvy et al., 2014).

Populations of *P. pristis* have undergone significant decline, primarily due to overfishing (especially as bycatch), habitat loss and illegal capture to remove the rostrum/teeth (Thorson, 1982; Cavanagh et al., 2005; Palmeira et al., 2013; Dulvy et al., 2016; Moore, 2017; Cabanillas-Torpoco et al., 2023). Australia remains one of the last refuges for four of the five existing sawfish species, making it a crucial region for conservation of these species (Morgan et al., 2017). In the Western Atlantic, the *P. pristis* population is believed to be restricted to coastal zones in some states of northern and northeastern Brazil, with occurrence of juveniles in the Amazon, Mearim River basins and Maranhão Gulf according to historical data (Charvet-Almeida and Faria 2008; Fernandez-Carvalho et al., 2014, Nunes et a., 2016, Feitosa et al., 2017). Although fishing for this species is prohibited under Brazilian law, there are still reports of capture by artisanal and industrial fishers (Schmid and Giarrizzo, 2017) in the region and the sale and consumption of *P. pristi*s meat continues, often under names that obscure the true origin of the product (Palmeira et al., 2013; Grant et al., 2021). Occurrence data for the Eastern Pacific region is even more uncertain. Records from 2014 and 2015 indicate its continued presence in the region in Peru, highlighting the need to identify and protect critical habitats that could contribute to sawfish conservation (Mendoza et al., 2017; Cabanillas-Torpoco et al., 2020; Espinoza et al., 2022).

## 5. Conclusion

The worldwide decline in the sawfishes follows trends for many elasmobranchs, with the aggravation that they have a greater tendency for mortality as bycatch in artisanal coastal fisheries because their rostrum get snagged in nets and because their rostra are also considered high value items in the illegal market. We have shown that because species-specific monitoring methods using eDNA have been developed based on genetic knowledge from a limited geographic region, this has limited the generalized use in other regions. The new mitogenomes and species-specific ddPCR primers and probe set developed here may contribute substantially to improving the potential to map and monitor these critically endangered species across the globe.

## Supporting information

All supplementary figures, tables and files

## Authors contributions

AESR and RMSB should be considered co-1st authors. Alan Érik S. Rodrigues: Data Curation; Formal Analysis; Investigation; Writing – original draft (lead); Writing – review and editing (equal). Rafaela Maria S. Brito: Formal analysis; Investigation; Writing – original draft; Writing – review and editing (equal). Patricia Charvet: Investigation; Resources; Writing – review and editing (equal). Vicente V. Faria: Investigation; Resources; Writing – review and editing (equal). Mariano Cabanillas-Torpoco: Investigation; Writing – review and editing (equal). Alexandre P. Aleixo: Funding Acquisition; Resources; Writing – review and editing (equal). Tibério César T. Burlamaqui: Investigation; Writing – original draft; Writing – review and editing (equal). Luis Fernando da S. Rodrigues-Filho: Writing – original draft; Writing – review and editing (equal). Angelico Asenjo: Investigation; Writing – review and editing (equal). Raquel Siccha-Ramirez: Investigation; Resources; Writing – review and editing (equal). Jorge Luiz S. Nunes: Investigation; Supervision; Writing – review and editing (equal). Hugo J. de Boer: Funding Acquisition; Resources; Writing – review and editing (equal). José Cerca: Writing – original draft (supporting); Writing – review and editing (equal). Quentin Mauvisseau: Conceptualization; Formal Analysis; Methodology; Project Administration; Resources; Supervision; Writing – original draft (supporting); Writing – review and editing (equal). Jonathan S. Ready: Conceptualization; Funding Acquisition; Investigation; Project Administration; Resources; Supervision; Writing – original draft (supporting); Writing – review and editing (equal). João Bráullio L. Sales: Conceptualization; Investigation; Project Administration; Resources; Supervision; Writing – original draft (supporting); Writing – review and editing (equal).

## Declaration of Generative AI and AI-assisted technologies

No generative AI or AI-assisted technology was used at any stage of this work.

## Supplementary information

Supplementary data to this article can be found online at (link to the page of the article)

## Data Availability Statement

All the Sequence Read Archive (SRA) required to replicate this study has been deposited in a BioProject on NLM-NCBI (PRJNA1217631).

## Acknowledgements

AESR thanks Bianca L. Paiva and Kamila de Fatima Silva for help with lab procedures. We thank all anonymous reviewers for comments on the manuscript.

## References

Andrews, S. 2010. FASTQC. A quality control tool for high throughput sequence data. Available online at: https://www.bioinformatics.babraham.ac.uk/projects/fastqc/

Anisimova, M., Gil, M., Dufayard, J.-F., Dessimoz, C., & Gascuel, O. (2011). Survey of Branch Support Methods Demonstrates Accuracy, Power, and Robustness of Fast Likelihood-based Approximation Schemes. Systematic Biology, 60(5), 685–699. 10.1093/sysbio/syr041

Bigelow, H.B. & Schroeder, W.C. (1953). Sharks, sawfishes, guitarfishes, skates and rays. Chimaeroids. In J. Tee–Van, C.M. Breder, S.F. Hildebrand, A.E. Parr, and W.C. Schroeder (Eds.), Fishes of the Western North Atlantic. Part 2. Sears Foundation for Marine Research, Yale University, New Haven (pp. 1–514).

Bleeker, P. (1851). Vijfde bijdrage tot de kennis der ichthyologische fauna van Borneo, met beschrijving van eenige nieuwe soorten van zoetwatervisschen. Natuurkundig Tijdschrift voor Nederlandsch Indië v. 2 (no. 3), 415–442.

Bohmann, K., Evans, A., Gilbert, M. T. P., Carvalho, G. R., Creer, S., Knapp, M., Yu, D. W., & de Bruyn, M. (2014). Environmental DNA for wildlife biology and biodiversity monitoring. Trends in Ecology & Evolution, 29(6), 358–367. 10.1016/j.tree.2014.04.003

Bommerlund, J., Baars, J.-R., Schrøder-Nielsen, A., Brys, R., Mauvisseau, C., de Boer, H., & Mauvisseau, Q. (2023). eDNA-based detection as an early warning tool for detecting established and emerging invasive amphipods. Management of Biological Invasions, 14(2), 321–333. 10.3391/mbi.2023.14.2.09

Bory de Saint-Vincent, J. B. G. M. (1822-31). Pisces accounts. In: Dictionnaire Classique d’Histoire Naturelle. Vols. 1-17. Paris (J. Tastu). Dictionnaire Classique d’Histoire Naturelle.

Bouckaert, R., Vaughan, T. G., Barido-Sottani, J., Duchêne, S., Fourment, M., Gavryushkina, A., Heled, J., Jones, G., Kühnert, D., De Maio, N., Matschiner, M., Mendes, F. K., Müller, N. F., Ogilvie, H. A., du Plessis, L., Popinga, A., Rambaut, A., Rasmussen, D., Siveroni, I., … Drummond, A. J. (2019). BEAST 2.5: An advanced software platform for Bayesian evolutionary analysis. PLOS Computational Biology, 15(4), e1006650. 10.1371/journal.pcbi.1006650

Boussarie, G., Bakker, J., Wangensteen, O. S., Mariani, S., Bonnin, L., Juhel, J.-B., Kiszka, J. J., Kulbicki, M., Manel, S., Robbins, W. D., Vigliola, L., & Mouillot, D. (2018). Environmental DNA illuminates the dark diversity of sharks. Science Advances, 4(5). 10.1126/sciadv.aap9661

Brys, R., Halfmaerten, D., Neyrinck, S., Mauvisseau, Q., Auwerx, J., Sweet, M., & Mergeay, J. (2021). Reliable eDNA detection and quantification of the European weather loach (*Misgurnus fossilis*). Journal of Fish Biology, 98(2), 399–414. 10.1111/jfb.14315

Budd, A. M., Cooper, M. K., Port, A. Le, Schils, T., Mills, M. S., Deinhart, M. E., Huerlimann, R., & Strugnell, J. M. (2021). First detection of critically endangered scalloped hammerhead sharks (*Sphyrna lewini*) in Guam, Micronesia, in five decades using environmental DNA. Ecological Indicators, 127, 107649. 10.1016/j.ecolind.2021.107649

Cabanillas-Torpoco, M., Castillo, D., Siccha-Ramírez, R., Forsberg, K., Purizaca, W., & Maceda, M. (2020). Occurrence of the largetooth sawfish Pristis pristis (Linneaus, 1758) in northern Peru. *Zootaxa*, 4868(1). 10.11646/zootaxa.4868.1.10

Cabanillas-Torpoco, M., Forsberg, K., Rosas-Luis, R., Bustamante Rosell, M., Ampuero-Portocarrero, C., Hernando, Á., Panizo, G., & Leeney, R. (2023). Status of the largetooth sawfish in Ecuador and Peru, and use of rostral teeth in cockfighting. Endangered Species Research, 52, 247–264. 10.3354/esr01279

Carlson, J., Blanco-Parra, MP, Bonfil-Sanders, R., Charles, R., Charvet, P., Chevis, M., Dulvy, N.K., Espinoza, M., Faria, V., Ferretti, F., Fordham, S., Giovos, I., Graham, J., Grubbs, D., Pacoureau, N. &amp; Phillips, N.M. (2022). Pristis pectinata. IUCN Red List of Threatened Species. 10.2305/IUCN.UK.2022-2.RLTS.T18175A58298676.en Accessed on 29 May 2024.

Castresana, J. (2000). Selection of Conserved Blocks from Multiple Alignments for Their Use in Phylogenetic Analysis. Molecular Biology and Evolution, 17(4), 540–552. 10.1093/oxfordjournals.molbev.a026334

Cavanagh, R. D., Camhi, M., Burgess, G. H., Cailliet, G. M., Fordham, S. V., Simpfendorfer, C. A., & Musick, J. A. (2005). *Sharks, rays and chimaeras: the status of the chondrichthyan fishes* (S. L. Folwer, Ed.). IUCN. 10.2305/IUCN.CH.2005.SSC-AP.9.en

Chan, P. P., & Lowe, T. M. (2019). tRNAscan-SE: Searching for tRNA Genes in Genomic Sequences. 1–14. 10.1007/978-1-4939-9173-0_1

Cooper, M. K., Huerlimann, R., Edmunds, R. C., Budd, A. M., Le Port, A., Kyne, P. M., Jerry, D. R., & Simpfendorfer, C. A. (2021). Improved detection sensitivity using an optimal eDNA preservation and extraction workflow and its application to threatened sawfishes. Aquatic Conservation: Marine and Freshwater Ecosystems, 31(8), 2131–2148. 10.1002/aqc.3591

Cooper, M. K., Villacorta□Rath, C., Burrows, D., Jerry, D. R., Carr, L., Barnett, A., Huveneers, C., & Simpfendorfer, C. A. (2022). Practical eDNA sampling methods inferred from particle size distribution and comparison of capture techniques for a Critically Endangered elasmobranch. Environmental DNA, 4(5), 1011–1023. 10.1002/edn3.279

Crampton-Platt, A., Yu, D. W., Zhou, X., & Vogler, A. P. (2016). Mitochondrial metagenomics: letting the genes out of the bottle. GigaScience, 5(1), 15. 10.1186/s13742-016-0120-y

Cuvier, G. (1829). Le Règne Animal, distribué d’après son organisation, pour servir de base à l’histoire naturelle des animaux et d’introduction à l’anatomie comparée. Edition 2. v. 2, i-xv + 1-406

de la Hoz Schilling, C., Jabado, R.W., Veríssimo, A., Caminiti, L., Sidina, E., Gandega, C. Y. Serrão, E. A. 2024. eDNA metabarcoding reveals a rich but threatened and declining elasmobranch community in West Africa’s largest marine protected area, the Banc d’Arguin. Conserv Genet (25), 805–821. 10.1007/s10592-024-01604-y

Dierckxsens, N., Mardulyn, P., & Smits, G. (2016). NOVOPlasty: *de novo* assembly of organelle genomes from whole genome data. Nucleic Acids Research, gkw955. 10.1093/nar/gkw955

Donath, A., Jühling, F., Al-Arab, M., Bernhart, S. H., Reinhardt, F., Stadler, P. F., Middendorf, M., & Bernt, M. (2019). Improved annotation of protein-coding genes boundaries in metazoan mitochondrial genomes. Nucleic Acids Research, 47(20), 10543– 10552. 10.1093/nar/gkz833

Doyle, J.J. & Doyle, J.L. (1987). A Rapid DNA Isolation Procedure for Small Quantities of Fresh Leaf Tissue. Phytochemical Bulletin, 19, 11–15.

Drummond, A. J., & Rambaut, A. (2007). BEAST: Bayesian evolutionary analysis by sampling trees. BMC Evolutionary Biology, 7(1), 214. 10.1186/1471-2148-7-214

Drummond, A. J., Ho, S. Y. W., Phillips, M. J., & Rambaut, A. (2006). Relaxed Phylogenetics and Dating with Confidence. PLoS Biology, 4(5), e88. 10.1371/journal.pbio.0040088

Dulvy, N. K., Davidson, L. N. K., Kyne, P. M., Simpfendorfer, C. A., Harrison, L. R., Carlson, J. K., & Fordham, S. V. (2016). Ghosts of the coast: global extinction risk and conservation of sawfishes. Aquatic Conservation: Marine and Freshwater Ecosystems, 26(1), 134–153. 10.1002/aqc.2525

Dulvy, N. K., Fowler, S. L., Musick, J. A., Cavanagh, R. D., Kyne, P. M., Harrison, L. R., Carlson, J. K., Davidson, L. N., Fordham, S. V, Francis, M. P., Pollock, C. M., Simpfendorfer, C. A., Burgess, G. H., Carpenter, K. E., Compagno, L. J., Ebert, D. A., Gibson, C., Heupel, M. R., Livingstone, S. R., … White, W. T. (2014). Extinction risk and conservation of the world’s sharks and rays. ELife, 3. 10.7554/eLife.00590

Edgar, R. C. (2004). MUSCLE: multiple sequence alignment with high accuracy and high throughput. Nucleic Acids Research, 32(5), 1792–1797. 10.1093/nar/gkh340

Espinoza, M., Bonfil-Sanders, R., Carlson, J., Charvet, P., Chevis, M., Dulvy, N.K., Everett, B., Faria, V., Ferretti, F., Fordham, S., Grant, M.I., Haque, A.B., Harry, A.V., Jabado, R.W., Jones, G.C.A., Kelez, S., Lear, K.O., Morgan, D.L., Phillips, N.M. &amp; Wueringer, B.E. (2022). “Pristis pristis”. IUCN Red List of Threatened Species. http://doi.org/10.2305/IUCN.UK.2022-2.RLTS.T18584848A58336780.en. Accessed on 29 May 2024.

Faria, V. V., Charvet-Almeida, P. (2008). Pristis pectinata. In: A.B.M. Machado; G.M. Drummond; A.P. Paglia. (Org.). Livro Vermelho da Fauna Brasileira Ameaçada de Extinção (Série Biodiversidade). 1ed.Belo Horizonte: Fundação Biodiversitas. v. II, p. 31–33

Faria, V. V., McDavitt, M. T., Charvet, P., Wiley, T. R., Simpfendorfer, C. A., & Naylor, G. J. P. (2013). Species delineation and global population structure of Critically Endangered sawfishes (Pristidae). Zoological Journal of the Linnean Society, 167(1), 136–164. 10.1111/j.1096-3642.2012.00872.x

Fee, K., Zabransky, K., Burgess, E., & Baeza, J. A. (2025). The complete mitochondrial genome of the imperiled Bullnose ray *Myliobatis freminvillei* (Myliobatiformes: Myliobatidae) with comments on its phylogenetic position and claims of diversifying selection affecting protein coding genes in a closely related species. Gene, 933, 148902. 10.1016/j.gene.2024.148902

Feitosa, L., Martins, A. B., & Nunes, J. S. (2017). Sawfish (Pristidae) records along the Eastern Amazon coast. Endangered Species Research, 34, 229–234. 10.3354/esr00852

Fernandez□Carvalho, J., Imhoff, J. L., Faria, V. V., Carlson, J. K., & Burgess, G. H. (2014). Status and the potential for extinction of the largetooth sawfish *Pristis pristis* in the Atlantic Ocean. Aquatic Conservation: Marine and Freshwater Ecosystems, 24(4), 478–497. 10.1002/aqc.2394

Feutry, P., Kyne, P., Pillans, R., Chen, X., Marthick, J., Morgan, D., & Grewe, P. (2015). Whole mitogenome sequencing refines population structure of the Critically Endangered sawfish *Pristis pristis*. Marine Ecology Progress Series, 533, 237–244. 10.3354/meps11354

Fordham SV, Jabado R, Kyne PM, Charvet P, Dulvy NK. 2018. *Saving Sawfish: Progress and Priorities*. IUCN Shark Specialist Group, Vancouver, Canada. 6 pp.

Gales, S. M., Parsons, K. T., Biesack, E. E., Ready, J., Siccha□Ramirez, R., Rosa, L. C., Rosa, R., Rotundo, M. M., Bills, R., Rodrigues, A. E. S., Rodrigues□Filho, L. F. S., McDowell, J., & Sales, J. B. L. (2024). Almost half of the *Gymnura* van Hasselt, 1823 species are unknown: Phylogeographic inference as scissors for cutting the hidden Gordian knot and clarify their conservation status. Journal of Systematics and Evolution, 62(4), 715– 738. 10.1111/jse.13027

Garman, S. (1906). New Plagiostomia. Bulletin of the Museum of Comparative Zoology v. 46 *(no.* *11*), 203–208.

Grant, J. R., Enns, E., Marinier, E., Mandal, A., Herman, E. K., Chen, C., Graham, M., Van Domselaar, G., & Stothard, P. (2023). Proksee: in-depth characterization and visualization of bacterial genomes. Nucleic Acids Research, 51(W1), W484–W492. 10.1093/nar/gkad326

Grant, M. I., White, W. T., Amepou, Y., Baje, L., Diedrich, A., Ibana, D., Jogo, D. J., Jogo, S., Kyne, P. M., Li, O., Mana, R., Mapmani, N., Nagul, A., Roeger, D., Simpfendorfer, C. A., & Chin, A. (2021). Local knowledge surveys with small□scale fishers indicate challenges to sawfish conservation in southern Papua New Guinea. Aquatic Conservation: Marine and Freshwater Ecosystems, 31(10), 2883–2900. 10.1002/aqc.3678

Grant, M.I., Charles, R., Fordham, S., Harry, A.V., Lear, K.O., Morgan, D.L., Phillips, N.M., Simeon, B., Wakhida, Y. &amp; Wueringer, B.E. (2022). Pristis clavata. In IUCN Red List of Threatened Species. 10.2305/IUCN.UK.2022-2.RLTS.T39390A68641215.en Accessed on 29 May 2024.

Hamilton, F. (1822). An account of the fishes found in the river Ganges and its branches. Edinburgh & London. i-vii + 1-405, Pls. 1-39.

Haque, A.B., Charles, R., D’Anastasi, B., Dulvy, N.K., Faria, V., Fordham, S., Grant, M.I., Harry, A.V., Jabado, R.W., Lear, K.O., Morgan, D.L., Tanna, A., Wakhida, Y. &amp; Wueringer, B.E. (2022). Anoxypristis cuspidata. In IUCN Red List of Threatened Species. http://doi.org/10.2305/IUCN.UK.2023-1.RLTS.T39389A58304073.en. Accessed on 29 May 2024.

Harrison, L.R. and Dulvy, N.K. (eds). 2014. Sawfish: A Global Strategy for Conservation. IUCN Species Survival Commission’s Shark Specialist Group, Vancouver, Canada.

Harry, A.V., Everett, B., Faria, V., Fordham, S., Grant, M.I., Haque, A.B., Ho, H., Jabado, R.W., Jones, G.C.A., Lear, K.O., Morgan, D.L., Phillips, N.M., Spaet, J.L.Y., Tanna, A. &amp; Wueringer, B.E. (2022). Pristis zijsron. In IUCN Red List of Threatened Species. 10.2305/IUCN.UK.2022-2.RLTS.T39393A58304631.en

Jordan, D. S. (1895). The fishes of Sinaloa. Proceedings of the California Academy of Sciences (Series 2) v. 5, 377–514, Pls. 26-55.

Kalyaanamoorthy, S., Minh, B. Q., Wong, T. K. F., von Haeseler, A., & Jermiin, L. S. (2017). ModelFinder: fast model selection for accurate phylogenetic estimates. Nature Methods, 14(6), 587–589. 10.1038/nmeth.4285

Kearse, M., Moir, R., Wilson, A., Stones-Havas, S., Cheung, M., Sturrock, S., Buxton, S., Cooper, A., Markowitz, S., Duran, C., Thierer, T., Ashton, B., Meintjes, P., & Drummond, A. (2012). Geneious Basic: An integrated and extendable desktop software platform for the organization and analysis of sequence data. Bioinformatics, 28(12), 1647–1649. 10.1093/bioinformatics/bts199

Kousteni, V., Mazzoleni, S., Vasileiadou, K., & Rovatsos, M. (2021). Complete Mitochondrial DNA Genome of Nine Species of Sharks and Rays and Their Phylogenetic Placement among Modern Elasmobranchs. Genes, 12(3), 324. 10.3390/genes12030324

Kumar, S., Stecher, G., Li, M., Knyaz, C., & Tamura, K. (2018). MEGA X: Molecular Evolutionary Genetics Analysis across Computing Platforms. Molecular Biology and Evolution, 35(6), 1547–1549. 10.1093/molbev/msy096

Kyne, P. M., Wang, J.-J., Xiang, D., Chen, X., & Feutry, P. (2018). The phylogenomic position of the Critically Endangered Largetooth Sawfish *Pristis pristis* (Rhinopristiformes, Pristidae), inferred from the complete mitochondrial genome. Mitochondrial DNA Part B, 3(2), 970–971. 10.1080/23802359.2018.1501315

Kyne, P., Oetinger, M., Grant, M., & Feutry, P. (2021). Life history of the Critically Endangered largetooth sawfish: a compilation of data for population assessment and demographic modelling. Endangered Species Research, 44, 79–88. 10.3354/esr01090

Lanfear, R., Frandsen, P. B., Wright, A. M., Senfeld, T., & Calcott, B. (2016). PartitionFinder 2: New Methods for Selecting Partitioned Models of Evolution for Molecular and Morphological Phylogenetic Analyses. *Molecular Biology and Evolution*, msw260. 10.1093/molbev/msw260

Last, P. R., Naylor, G. J. P., & Manjaji-Matsumoto, B. M. (2016). A revised classification of the family Dasyatidae (Chondrichthyes: Myliobatiformes) based on new morphological and molecular insights. Zootaxa, 4139(3). 10.11646/zootaxa.4139.3.2

Latham, J. F. (1794). An essay on the various species of sawfish. The Transactions of the Linnean Society of London v. 2 *(art. 25)*, 273–282, Pls. 26-27.

Leeney, R. H., & Downing, N. (2016). Sawfishes in The Gambia and Senegal – shifting baselines over 40□years. Aquatic Conservation: Marine and Freshwater Ecosystems, 26(2), 265–278. 10.1002/aqc.2545

Lehman, R. N., Poulakis, G. R., Scharer, R. M., Schweiss, K. E., Hendon, J. M., & Phillips, N. M. (2020). An environmental DNA tool for monitoring the status of the Critically Endangered Smalltooth Sawfish, *Pristis pectinata*, in the western Atlantic. Conservation Genetics Resources, 12(4), 621–629. 10.1007/s12686-020-01149-5

Linnaeus [C.] (1758). Systema Naturae, Ed. X. (Systema naturae per regna tria naturae, secundum classes, ordines, genera, species, cum characteribus, differentiis, synonymis, locis. Tomus I. Editio decima, reformata.) Holmiae. v. 1: i-ii + 1-824 [Nantes and Pisces in Tom. 1, pp. 230–338;

Love, M. S., Bizzarro, J. J., Cornthwaite, A. M., Frable, B. W., & Maslenikov, K. P. (2021). Checklist of marine and estuarine fishes from the Alaska–Yukon Border, Beaufort Sea, to Cabo San Lucas, Mexico. Zootaxa, 5053(1), 1–285. 10.11646/zootaxa.5053.1.1

Mauvisseau, Q., Halfmaerten, D., Neyrinck, S., Burian, A., & Brys, R. (2021). Effects of preservation strategies on environmental DNA detection and quantification using ddPCR. Environmental DNA, 3(4), 815–822. 10.1002/edn3.188

McDavitt, M. T. (2014). Sawfish products and trade. In: Harrison LR, Dulvy NK (Eds) Sawfish: A Global Strategy for Conservation. International Union for the Conservation of Nature Species Survival Commission’s Shark Specialist Group, Vancouver, 72–75.

Mendoza, A., Kelez, S., Cherres, W. G., & Maguiño, R. (2017). The Largetooth Sawfish, Pristis pristis (Linnaeus, 1758), is not extirpated from Peru: new records from Tumbes. Check List, 13(4), 261–265. 10.15560/13.4.261

Minh, B. Q., Nguyen, M. A. T., & von Haeseler, A. (2013). Ultrafast Approximation for Phylogenetic Bootstrap. Molecular Biology and Evolution, 30(5), 1188–1195. 10.1093/molbev/mst024

Minh, B. Q., Schmidt, H. A., Chernomor, O., Schrempf, D., Woodhams, M. D., von Haeseler, A., & Lanfear, R. (2020). IQ-TREE 2: New Models and Efficient Methods for Phylogenetic Inference in the Genomic Era. Molecular Biology and Evolution, 37(5), 1530– 1534. 10.1093/molbev/msaa015

Miya, M., & Nishida, M. (1999). Organization of the Mitochondrial Genome of a Deep-Sea Fish, Gonostoma gracile (Teleostei: Stomiiformes): First Example of Transfer RNA Gene Rearrangements in Bony Fishes. Marine Biotechnology, 1(5), 416–426. 10.1007/PL00011798

Miya, M., & Nishida, M. (2015). The mitogenomic contributions to molecular phylogenetics and evolution of fishes: a 15-year retrospect. Ichthyological Research, 62(1), 29–71. 10.1007/s10228-014-0440-9

Moore, A. (2017). Are guitarfishes the next sawfishes? Extinction risk and an urgent call for conservation action. Endangered Species Research, 34, 75–88. 10.3354/esr00830

Moore, A. B. M. (2015). A review of sawfishes (Pristidae) in the Arabian region: diversity, distribution, and functional extinction of large and historically abundant marine vertebrates. Aquatic Conservation: Marine and Freshwater Ecosystems, 25(5), 656–677. 10.1002/aqc.2441

Morgan, D., Ebner, B., Allen, M., Gleiss, A., Beatty, S., & Whitty, J. (2017). Habitat use and site fidelity of neonate and juvenile green sawfish *Pristis zijsron* in a nursery area in Western Australia. Endangered Species Research, 34, 235–249. 10.3354/esr00847

Näsvall, K., Boman, J., Talla, V., & Backström, N. (2023). Base Composition, Codon Usage, and Patterns of Gene Sequence Evolution in Butterflies. Genome Biology and Evolution, 15(8). 10.1093/gbe/evad150

Naylor, G. J., Caira, J. N., Jensen, K., Rosana, K. A., Straube, N., & Lakner, C. (2012). Elasmobranch phylogeny: a mitochondrial estimate based on 595 species. Biology of sharks and their relatives, 2, 31–56.

Nunes, J. L. S., Rincon, G., Piorski, N. M., & Martins, A. P. B. (2016). Near□term embryos in a *Pristis pristis* (Elasmobranchii: Pristidae) from Brazil. Journal of Fish Biology, 89(1), 1112–1120. 10.1111/jfb.12946

Palmeira, C. A. M., da Silva Rodrigues-Filho, L. F., de Luna Sales, J. B., Vallinoto, M., Schneider, H., & Sampaio, I. (2013). Commercialization of a critically endangered species (largetooth sawfish, Pristis perotteti) in fish markets of northern Brazil: Authenticity by DNA analysis. Food Control, 34(1), 249–252. 10.1016/j.foodcont.2013.04.017

Petean, F. F., Yang, L., Corrigan, S., Lima, S. M. Q., & Naylor, G. J. P. (2024). How many linaeges are there of the stingrays genus *Hypanus* (Myliobatiformes: Dasyatidae) and why does it matter? Neotropical Ichthyology, 22(1). 10.1590/1982-0224-2023-0046

Phillips, N. M., Chaplin, J. A., Morgan, D. L., & Peverell, S. C. (2011). Population genetic structure and genetic diversity of three critically endangered *Pristis* sawfishes in Australian waters. Marine Biology, 158(4), 903–915. 10.1007/s00227-010-1617-z

Rambaut A. & Drummond A. (2009). Tracer v1. 5. Available from: http://tree.bio.ed.ac.uk/software/tracer/

Rambaut, A. (2018). Figtree ver 1.4.4. Institute of Evolutionary Biology, University of Edinburgh, Edinburgh.

Robillard, M., & Séret, B. (2006). Cultural importance and decline of sawfish (Pristidae) populations in West Africa. Cybium, 30(4), 23–30.

Rodrigues-Filho, L. F. S., Costa Nogueira, P., Sodré, D., Silva Leal, J. R., Nunes, J. L. S., Rincon, G., Lessa, R. P. T., Sampaio, I., Vallinoto, M., Ready, J. S., & Sales, J. B. L. (2023). Evolutionary History and Taxonomic Reclassification of the Critically Endangered Daggernose Shark, a Species Endemic to the Western Atlantic. Journal of Zoological Systematics and Evolutionary Research, 2023, 1–16. 10.1155/2023/4798805

Rosa, F. dos A. S., Gasalla, M. A., de Queiroz, A. K. O., Ribas, T. F. A., Mauvisseau, Q., de Boer, H. J., Thorbek, B. L. G., Oliveira, R. R. M., Laux, M., Postuma, F. A., & Ready, J. S. (2024). Molecular analyses of carangid fish diets reveal interLpredation, dietary overlap, and the importance of early life stages in trophic ecology. Ecology and Evolution, 14(1). 10.1002/ece3.10817

Rozas, J., Ferrer-Mata, A., Sánchez-DelBarrio, J. C., Guirao-Rico, S., Librado, P., Ramos-Onsins, S. E., & Sánchez-Gracia, A. (2017). DnaSP 6: DNA Sequence Polymorphism Analysis of Large Data Sets. Molecular Biology and Evolution, 34(12), 3299–3302. 10.1093/molbev/msx248

Rubinoff, D. (2006). Utility of Mitochondrial DNA Barcodes in Species Conservation. Conservation Biology, 20(4), 1026–1033. 10.1111/j.1523-1739.2006.00372.x

Sales, J. B. L., de Oliveira, C. N., dos Santos, W. C. R., Rotundo, M. M., Ferreira, Y., Ready, J., Sampaio, I., Oliveira, C., Cruz, V. P., Lara-Mendoza, R. E., & da Silva Rodrigues-Filho, L. F. (2019). Phylogeography of eagle rays of the genus *Aetobatus*: *Aetobatus narinari* is restricted to the continental western Atlantic Ocean. Hydrobiologia, 836(1), 169–183. 10.1007/s10750-019-3949-0

Schmid, K., & Giarrizzo, T. (2017). New record of a largetooth sawfish specimen from the Amazon River estuary in Northern Brazil. Fisheries, 42(*5*), 254–255. 10.1080/03632415.2017.1276357

Sharks, sawfishes, guitarfishes, skates and rays. Chimaeroids. In J. Tee–Van, C.M. Breder, S.F. Hildebrand, A.E. Parr, and W.C. Schroeder (Eds.), Fishes of the Western North Atlantic. Part 2. Sears Foundation for Marine Research, Yale University, New Haven (pp. 1–514)

Simpfendorfer, C., Kyne, P., Noble, T., Goldsbury, J., Basiita, R., Lindsay, R., Shields, A., Perry, C., & Jerry, D. (2016). Environmental DNA detects Critically Endangered largetooth sawfish in the wild. Endangered Species Research, 30, 109–116. 10.3354/esr00731

Taite, M., Fernández-Álvarez, F. Á., Braid, H. E., Bush, S. L., Bolstad, K., Drewery, J., S., Mills, J.M. Strugnell, M. Vecchione, R. Villanueva, J.R. & Allcock, A. L. (2023). Genome skimming elucidates the evolutionary history of Octopoda. Molecular Phylogenetics and Evolution, 182, 107729.

Thorson, T., B. (1982). The impact of commercial exploitation on sawfish and shark populations in Lake Nicaragua. Fisheries 7: 2−10

Wang, C., Ye, P., Pillans, R., Chen, X., Wang, J., & Feutry, P. (2023). Evolution of the Critically Endangered Green Sawfish *Pristis zijsron* (Rhinopristiformes, Pristidae), Inferred from the Whole Mitochondrial Genome. Genes, 14(11), 2052. 10.3390/genes14112052

West, K. M., Stat, M., Harvey, E. S., Skepper, C. L., DiBattista, J. D., Richards, Z. T., Travers, M. J., Newman, S. J., & Bunce, M. (2020). eDNA metabarcoding survey reveals fineLscale coral reef community variation across a remote, tropical island ecosystem. Molecular Ecology, 29(6), 1069–1086. 10.1111/mec.15382

White, W. T., & Naylor, G. J. P. (2016). Resurrection of the family Aetobatidae (Myliobatiformes) for the pelagic eagle rays, genus *Aetobatus*. Zootaxa, 4139(3). 10.11646/zootaxa.4139.3.10

White, W. T., Furumitsu, K., & Yamaguchi, A. (2013). A New Species of Eagle Ray Aetobatus narutobiei from the Northwest Pacific: An Example of the Critical Role Taxonomy Plays in Fisheries and Ecological Sciences. PLoS ONE, 8(12), e83785. 10.1371/journal.pone.0083785

Wickham, H. (2016). ggplot2: Elegant Graphics for Data Analysis. Springer-Verlag New York. ISBN 978-3-319-24277-4. https://ggplot2.tidyverse.org.

Wilcox, T. M., Carim, K. J., McKelvey, K. S., Young, M. K., & Schwartz, M. K. (2015). The Dual Challenges of Generality and Specificity When Developing Environmental DNA Markers for Species and Subspecies of *Oncorhynchus*. PLOS ONE, 10(11), e0142008. 10.1371/journal.pone.0142008

Winn, J. C., Maduna, S. N., & Bester-van der Merwe, A. E. (2024). A comprehensive phylogenomic study unveils evolutionary patterns and challenges in the mitochondrial genomes of Carcharhiniformes: A focus on Triakidae. Genomics, 116(1), 110771. 10.1016/j.ygeno.2023.110771

Xiang, C., Gao, F., Jakovlić, I., Lei, H., Hu, Y., Zhang, H., Zou, H., Wang, G., & Zhang, D. (2023). Using PhyloSuite for molecular phylogeny and treeLbased analyses. IMeta, 2(1). 10.1002/imt2.87

Yokobori, S., Fukuda, N., Nakamura, M., Aoyama, T. & Oshima, T. 2004. Long-term conservation of six duplicated structural genes in cephalopod mitochondrial genomes. Mol. Biol. Evol. 21, 2034–2046. 10.1093/molbev/msh227

Zhang, J., Miao, G., Hu, S., Sun, Q., Ding, H., Ji, Z., Guo, P., Yan, S., Wang, C., Kan, X., & Nie, L. (2021). Quantification and evolution of mitochondrial genome rearrangement in Amphibians. BMC Ecology and Evolution, 21(1), 19. 10.1186/s12862-021-01755-3

Zhu, T., Sato, Y., Sado, T., Miya, M., & Iwasaki, W. (2023). MitoFish, MitoAnnotator, and MiFish Pipeline: Updates in 10 Years. Molecular Biology and Evolution, 40(3). 10.1093/molbev/msad035

