## Supplementary material for "Geographical variation in mitogenomes of the largetooth sawfish *Pristis pristis*: challenges and perspectives for conservation efforts": All supplementary figures, tables and files

Supporting Information

**Appendix A**: Sequences utilized for validation of the *in-silico* test with new primers/probes and eDNA samples metadata.

**Table A.1**. GenBank accession numbers of the targeted, closely related and/or potentially co-occurring species used during for the *in-silico* development and validation of the species-specific primers and probe developed in this study.

| | |
|---|---|
| MF977764; MH005928;MN105755; MN105758; MN105759; MN105760; MN105761; MN105762; MN105763; MN105764; MN105765; MN105766; MN105767; MN105840; MN105841; NC_039438 | *Pristis pristis* (Linnaeus, 1758) |
| KP400584; MF682494; NC_027182 | *Pristis pectinata* (Latham, 1794) |
| JN184072; KF381507; NC_022821 | *Pristis clavata* (Garman, 1906) |
| EU398989; EU784162; MH005927; MH825681 | *Pristis zijsron* (Bleeker, 1851) |
| OR284669; OR284668; OR227124; OR227123; OR227122; OR227121; OR227120; OR227119; OR227118; OR227117; OR227116; OR227115; OR227113; OR227111; OQ386120; OQ385079; OQ385078; OQ385077; OQ385076; OQ385075; OQ361661; OQ221446; OQ221402; OQ221401; OQ221400; OQ221399; OQ221398; OQ221397; OQ221396; OQ221395; OQ221394; OQ221393; OQ221392; OQ221391; OQ221390; OQ221389; OQ221388; OQ221387; OQ221386; OQ221385; OQ221384; OQ221383; OQ221382; OQ221381; OQ221380; OQ221379; OQ221378; OQ221377; OQ221376; OQ221375; OQ221374; OQ221373; OQ221372; OQ221371; OQ221370; OQ221369; OQ221368; OQ221367; OQ221366; OQ221365; OQ221364; OQ221363; OQ221362; OQ221361; OQ221360; OQ221359; OQ221357; OQ221356; OQ221355; OQ221354; OQ221353; OQ221352; OQ221351; OQ221350; OQ221349; OQ221348; OQ221347; OQ221346; OQ221345; OQ221344; OQ221343; OQ221342; OQ221341; OQ221340; OQ221339; OQ221338; OQ221337; OQ221336; OQ221335; OQ221334; OQ221333; OQ221332; OQ221331; OQ221330; OQ221329; OQ221328; OQ221327; OQ221326; OQ221325; OQ221324; OQ221323; OQ221322; OQ221321; OQ221320; OQ221319; OQ221318; OQ221317; OQ221316; OQ221315; OQ221314; OQ221313; OQ221312; OQ221311; OQ221310; OQ221309; OQ221308; OQ221307; OQ221306; OQ221305; OQ221304; OP800195; ON678608; ON678607; ON678606; ON678605; ON678604; ON678603; ON678602; ON678601; ON678600; ON678599; ON678598; | *Rhynchobatus australiae* (Whitley, 1939) (and other sequences referred as *Rhynchobatus cf. australiae*) |

| | |
|---|---|
| ON678597; ON678596; ON678595; ON678594; ON678593; ON678592; ON678591; ON678590; ON678589; ON678588; ON678587; ON678586; ON678585; ON678584; ON678583; ON678582; ON678581; ON678580; ON678579; ON678578; ON678577; ON678576; ON678575; ON678574; ON678573; ON678572; ON678571; ON678570; ON678569; ON678568; ON678567; ON678566; ON678565; ON678564; ON678563; ON678562; ON678561; ON678560; ON678559; ON678558; ON678557; ON678556; ON678555; NC_030254; MW514055; MW509730; MW509729; MW509728; MW509727; MW509726; MW509725; MW509724; MW509723; MW509722; MW509721; MW509720; MW509719; MW509718; MW509717; MW509716; MW509715; MW509714; MW509713; MW509712; MW509711; MW509710; MT983932; MT983931; MT983930; MT933190; MT933189; MT933188; MT933187; MT933186; MT002425; MN795536; MK422139; MH243231; MH243226; MH243224; MH243223; MH243213; MH243211; MH243210; MH243209; MH243207; MH243204; MH243203; MH243197; MH243195; MH243192; MH243189; MH243188; MH243169; MH243111; MH243109; MG792126; MG774928; MG774925; MG644272; MF508696; KU936207; KU746824; KU255184; KP719753; JN108019; JN108018; JN022596; JN022595; EU399009; EU399008; EU399007; DQ108199; GU674342 | |
| OQ221448; OQ221447; OQ221445; OQ221444; OQ221443; OQ221442; OQ221441; OQ221440; OQ221439; OQ221438; OQ221437; OQ221436; OQ221435; OQ221434; OQ221433; OQ221432; OQ221431; OQ221430; OQ221429; OQ221428; OQ221427; OQ221426; OQ221425; OQ221424; OQ221423; OQ221422; OQ221421; OQ221420; OQ221419; OQ221418; OQ221417; OQ221416; OQ221415; OQ221414; OQ221413; OQ221412; OQ221411; OQ221410; OQ221409; OQ221408; OQ221407; OQ221405; OQ221404; OQ221403; ON065568; NC_066688; JN184077; JF494386; JF494385; JF494384; GU805049 | *Rhynchobatus djiddensis* (Forsskål, 1775) |
| DQ108191; DQ108192; DQ108197; DQ108198; EU399010; KF899376; KF899377; KF899689; KF899690; KF899691; KF899692; KF899693; KJ825840; MH243146; MH243205; MH243229; MN988687; MW517841; NC_047241; ON678283; ON678284; ON678285 | *Rhynchobatus laevis* (Bloch & Schneider, 1801) (and other sequences referred as *Rhynchobatus cf. laevis*) |

**Table A.2**. GPS coordinates, sampling volume, site replicate number and sampling date of each eDNA sample analysed in this study.

| GPS coordinates | | Total sampling volume (L) | Replicate number | Sampling date | Other info |
|---|---|---|---|---|---|
| Lat | Long | | | | |
| -44.832.776 | -3.383.594 | 1.04 | 4 | 8/23/2021 | 0.8 µm filter pore size |
| -41.840.232 | -2.823.009 | 1.00 | 4 | 8/25/2021 | 0.8 µm filter pore size |
| -41.810.174 | -2.755.846 | 1.00 | 4 | 8/25/2021 | 0.8 µm filter pore size |
| -41.654.059 | -2.878.445 | 1.00 | 4 | 8/25/2021 | 0.8 µm filter pore size |
| -41.903.141 | -2.891.949 | 1.00 | 4 | 8/26/2021 | 0.8 µm filter pore size |
| -42.254.833 | -2.856.815 | 1.00 | 4 | 8/26/2021 | 0.8 µm filter pore size |
| -42.260.557 | -2.762.330 | 1.00 | 4 | 8/26/2021 | 0.8 µm filter pore size |
| -42.273.248 | -2.765.176 | 1.00 | 4 | 8/27/2021 | 0.8 µm filter pore size |
| -44.106.099 | -2.416.492 | 1.00 | 4 | 8/28/2021 | 0.8 µm filter pore size |
| -44.056.149 | -2.566.221 | 1.00 | 4 | 8/28/2021 | 0.8 µm filter pore size |
| -44.318.915 | -2.507.157 | 1.00 | 4 | 8/28/2021 | 0.8 µm filter pore size |
| -44.477.591 | -2.830.890 | 1.00 | 4 | 8/29/2021 | 0.8 µm filter pore size |
| -44.398.085 | -2.715.915 | 1.00 | 4 | 8/29/2021 | 0.8 µm filter pore size |
| -44.360.471 | -2.647.056 | 1.00 | 4 | 8/29/2021 | 0.8 µm filter pore size |
| -44.894.688 | -3.293.452 | 1.00 | 4 | 8/30/2021 | 0.8 µm filter pore size |
| -44.674.165 | -2.886.536 | 1.00 | 4 | 8/30/2021 | 0.8 µm filter pore size |
| -45.069.429 | -2.527.155 | 1.00 | 4 | 8/30/2021 | 0.8 µm filter pore size |
| -45.362.191 | -1.674.754 | 0.50 | 4 | 8/31/2021 | 0.8 µm filter pore size |

| | | | | | |
|---|---|---|---|---|---|
| -46.316.071 | -1.799.275 | 0.60 | 4 | 8/31/2021 | 0.8 µm filter pore size |
| -47.455.646 | -0.761001 | 0.40 | 4 | 9/1/2021 | 0.8 µm filter pore size |
| -46.761.968 | -1.056.744 | 0.86 | 4 | 10/2/2021 | 0.8 µm filter pore size |
| -47.367.586 | -0.624614 | 1.00 | 4 | 11/2/2021 | 0.8 µm filter pore size |
| -47.171.746 | -0.763368 | 1.00 | 4 | 11/2/2021 | 0.8 µm filter pore size |

*Table C2 supporting information:* eDNA sampling protocol

A total of 92 samples across 23 locations were sampled from west of Maranhão to northeast of Pará in Brazil (see Table C2). At each location, water samples consisted of subsamples collected from the surface. Samples from each location were then filtered using syringes through a sterile 0.8 µm filter membrane with a swinnex filter holder. Following filtration, filters were preserved using the Longmire's buffer before eDNA extraction. Extractions were completed in a PCR free room and performed using DNeasy PowerSoil Kit (QIAGEN) following the manufacturer's instructions and eluted in a final 50 µL volume. Sampled sites were selected as a part of another project and included in the testing workflow of the newly designed primers/probe due to the high species richness expected in these locations.

**Appendix B**: tRNA Structures of tRNAs from both specimens of *P. pristis*, Western Atlantic and Eastern Pacific.

#### Western Atlantic:

Your-seq.trna1 Phe (GAA) 87.0 bits

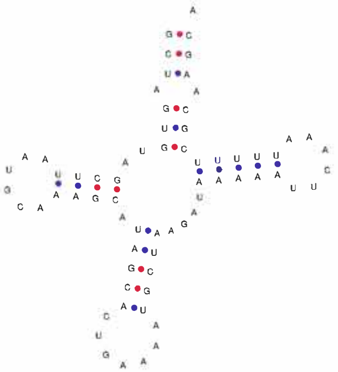

Your-seq.trna Val (TAC)

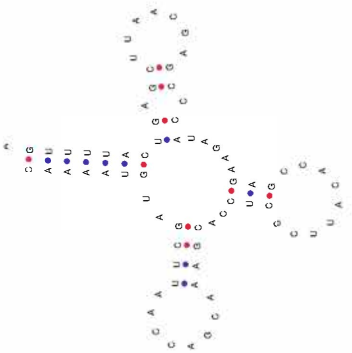

Your-seq.trna1 Leu (TAA) 117.8 bits

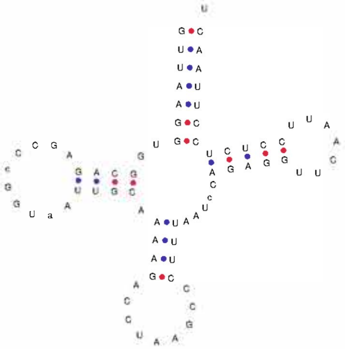

Your-seq.trna1 Ile (GAT) 91.5 bits

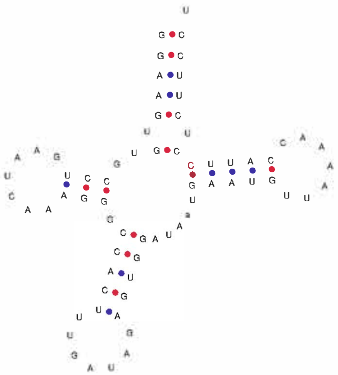

Your-seq.trna1 Gln (TTG) 100.8 bits

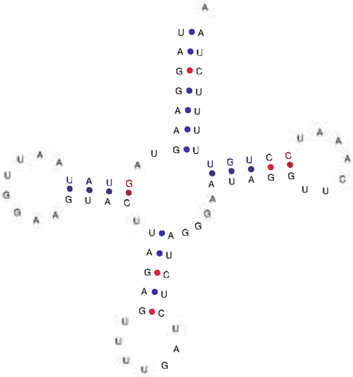

Your-seq.trna1 Met (CAT) 111.6 bits

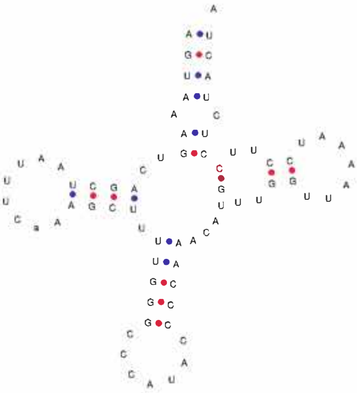

Your-seq.trna1 Trp (TCA) 96.2 bits

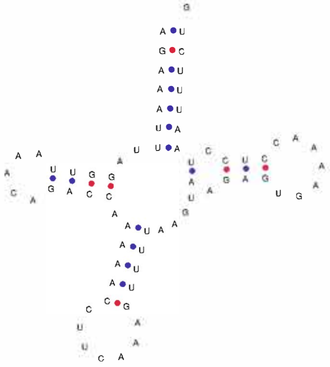

Your-seq.trna1 Ala (TGC) 92.1 bits

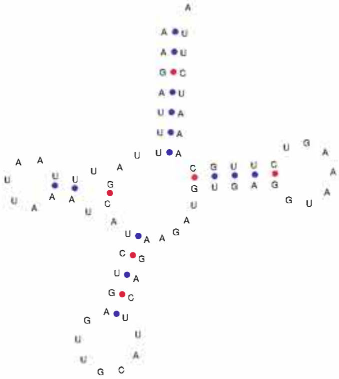

Your-seq.trna1 Asn (GTT) 96.3 bits

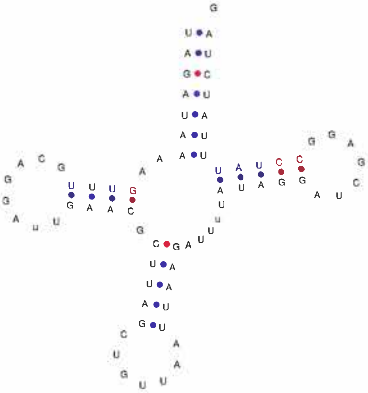

Your-seq.trna1 Cys (GCA) 77.6 bits

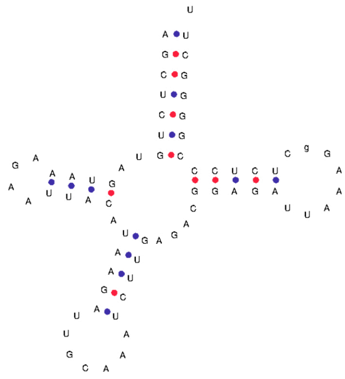

Your-seq.trna1 Tyr (GTA) 91.6 bits

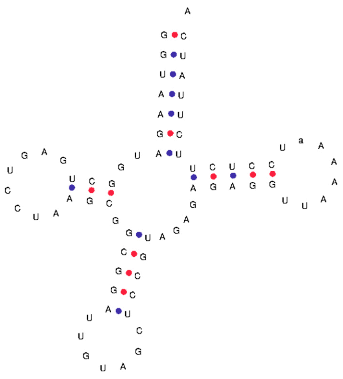

Your-seq.trna1 Ser (TGA) 112.1 bits

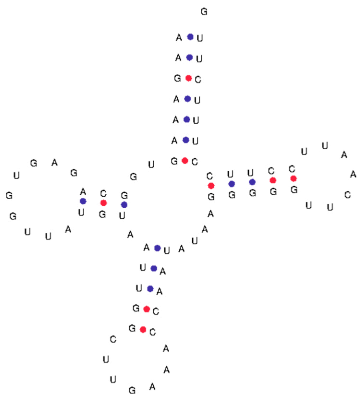

Your-seq.trna1 Asp (GTC) 89.3 bits

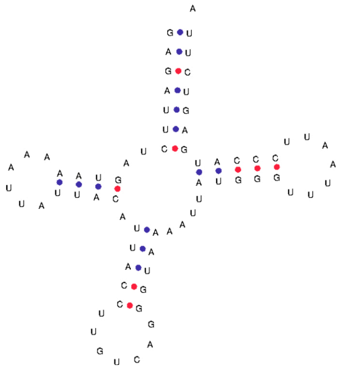

Your-seq.trna1 Lys (TTT) 109.3 bits

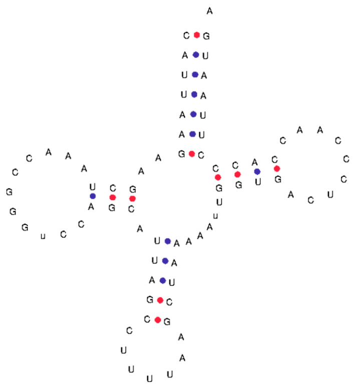

Your-seq.trna1 Gly (TCC) 96.7 bits

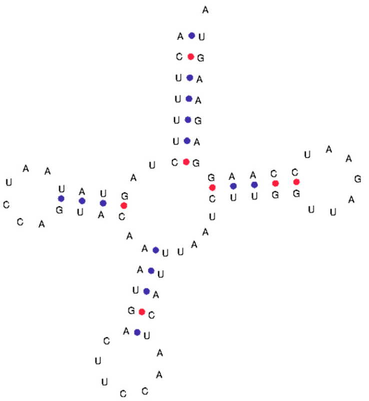

Your-seq.trna1 Arg (TCG) 85.6 bits

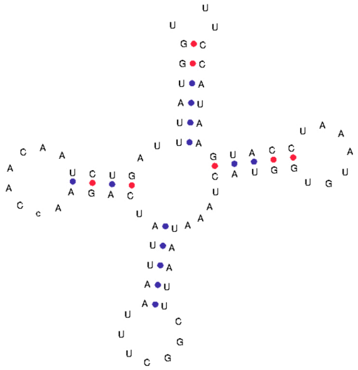

Your-seq.trna1 His (GTG) 101.3 bits

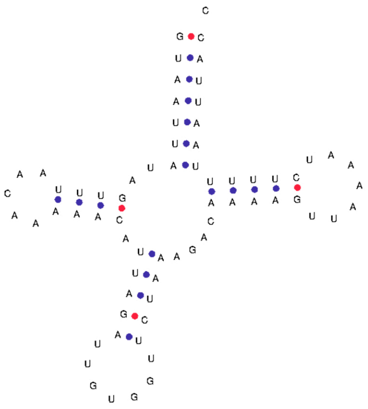

Your-seq.trna1 Ser (GCT) 59.3 bits

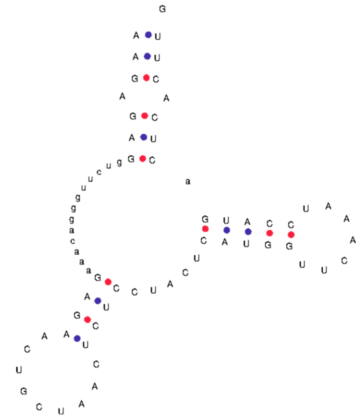

Your-seq.trna1 Leu (TAG) 115.0 bits

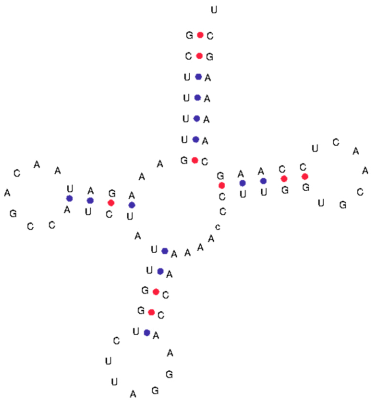

Your-seq.trna1 Glu (TTC) 94.7 bits

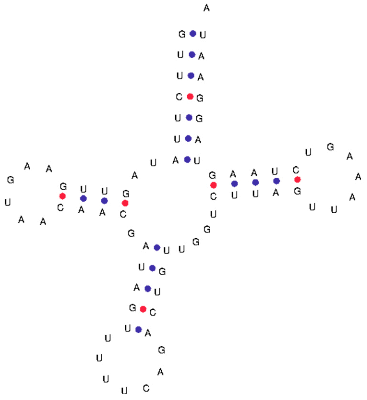

Your-seq.trna1 Thr (TGT) 78.2 bits

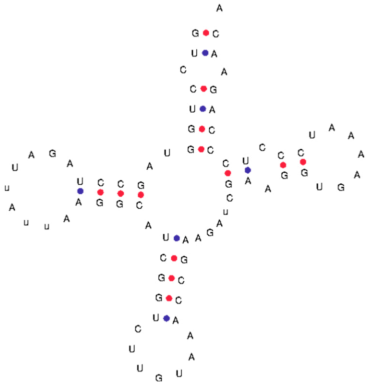

Your-seq.trna Pro (TGG)

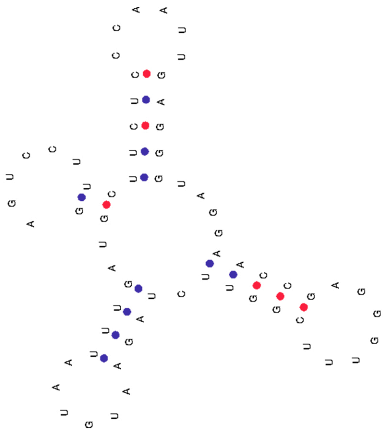

---

### Eastern Pacific:

Contig1.trna1 Phe (GAA) 87.0 bits

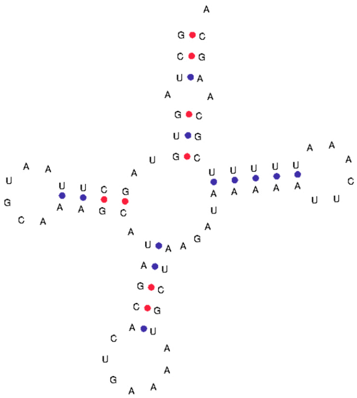

Your-seq.trna Val (TAC)

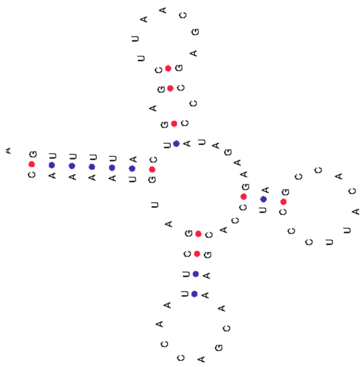

Contig1.trna3 Leu (TAA) 117.8 bits

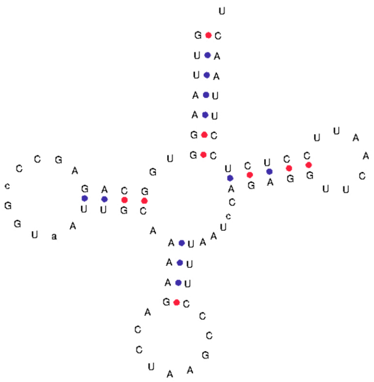

Contig1.trna4 Ile (GAT) 91.5 bits

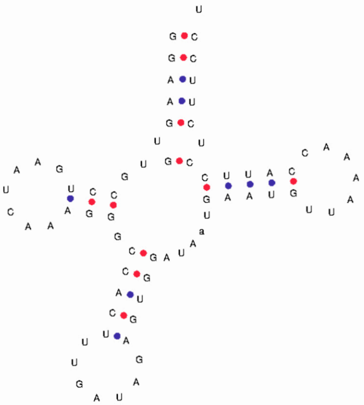

Contig1.trna23 Gln (TTG) 100.8 bits

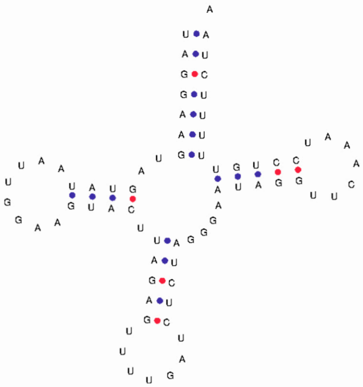

Contig1.trna5 Met (CAT) 109.5 bits

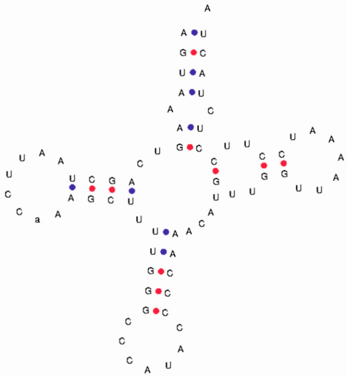

Contig1.trna6 Trp (TCA) 101.8 bits

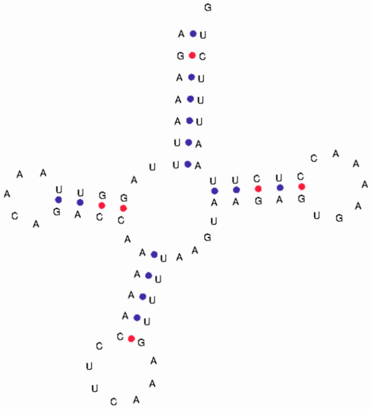

Contig1.trna22 Ala (TGC) 92.1 bits

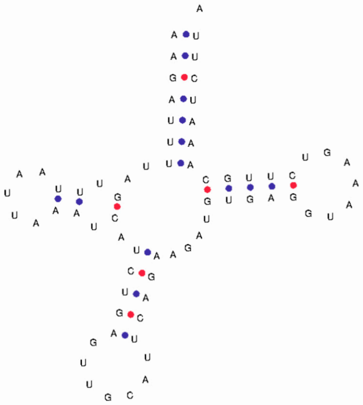

Contig1.trna21 Asn (GTT) 96.3 bits

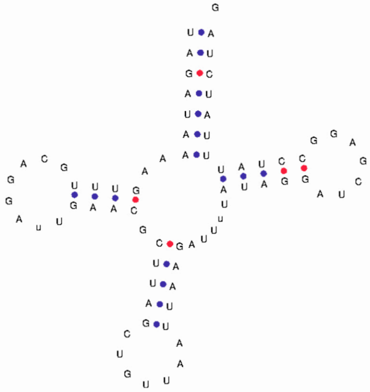

Contig1.trna20 Cys (GCA) 79.3 bits

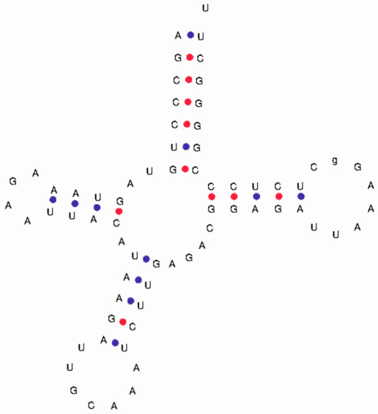

Contig1.trna19 Tyr (GTA) 94.6 bits

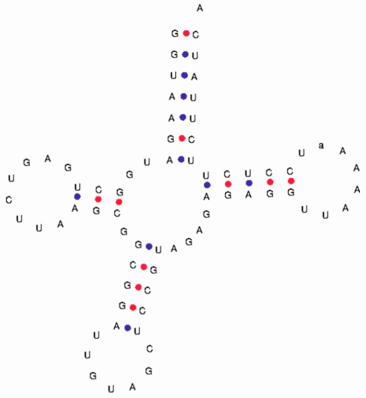

Contig1.trna18 Ser (TGA) 112.1 bits

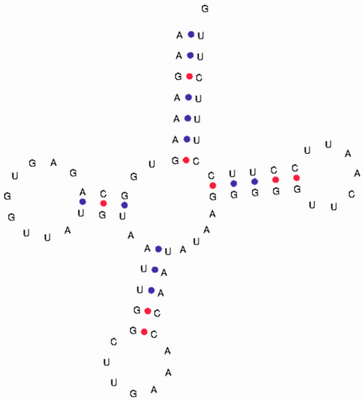

Contig1.trna7 Asp (GTC) 89.3 bits

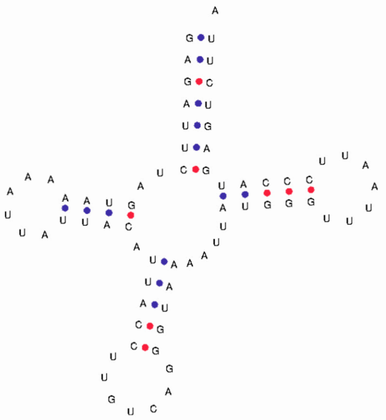

Contig1.trna8 Lys (TTT) 109.3 bits

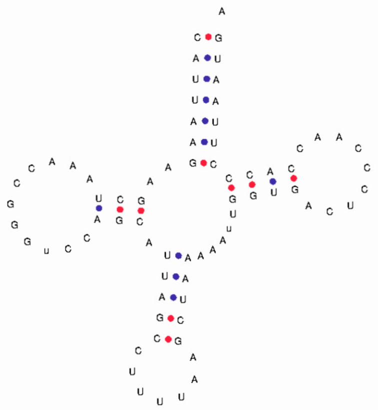

Contig1.trna9 Gly (TCC) 96.7 bits

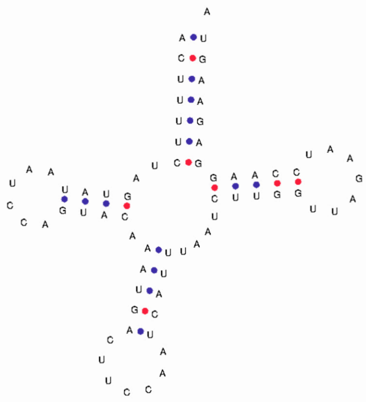

Contig1.trna10 Arg (TCG) 85.6 bits

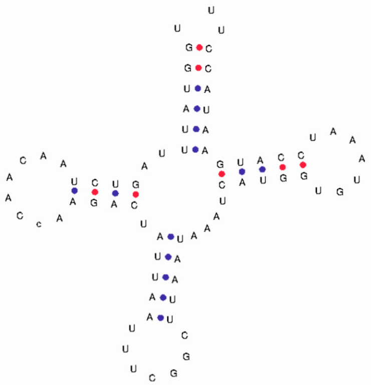

Contig1.trna11 His (GTG) 101.3 bits

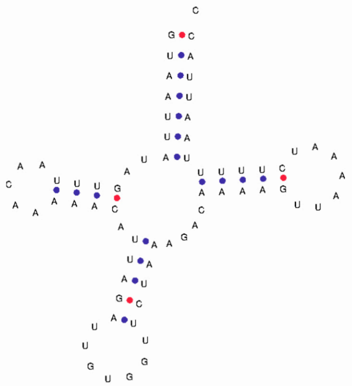

trna Ser (GCT)

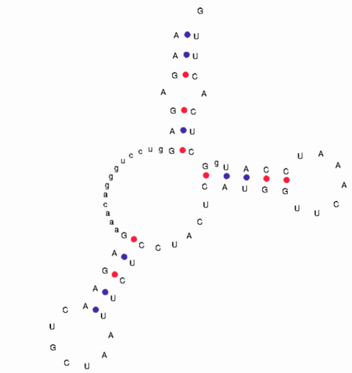

Contig1.trna13 Leu (TAG) 115.0 bits

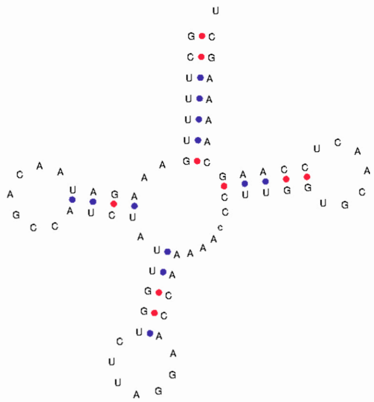

Contig1.trna17 Glu (TTC) 91.7 bits

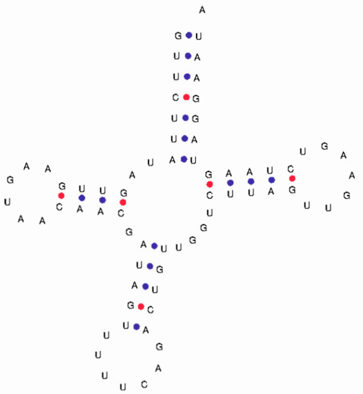

Contig1.trna14 Thr (TGT) 78.9 bits

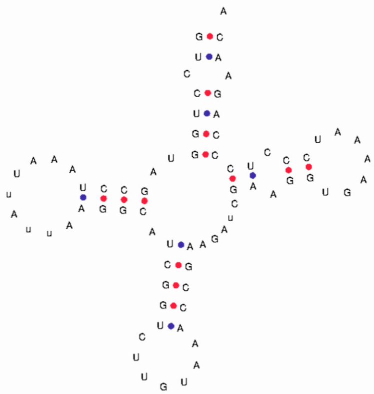

Contig1.trna16 Pro (TGG) 78.7 bits

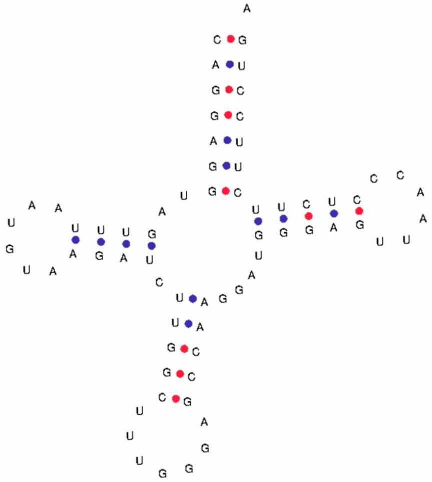

Contig1.trna15 Pro (TGG) 61.1 bits

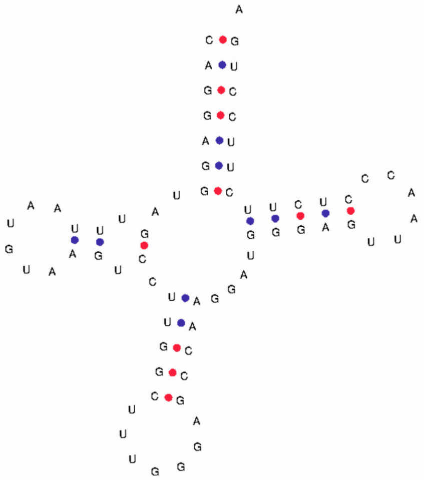

**Appendix C**: RSCU content, Nucleotide diversity and Phylogenetics analysis.

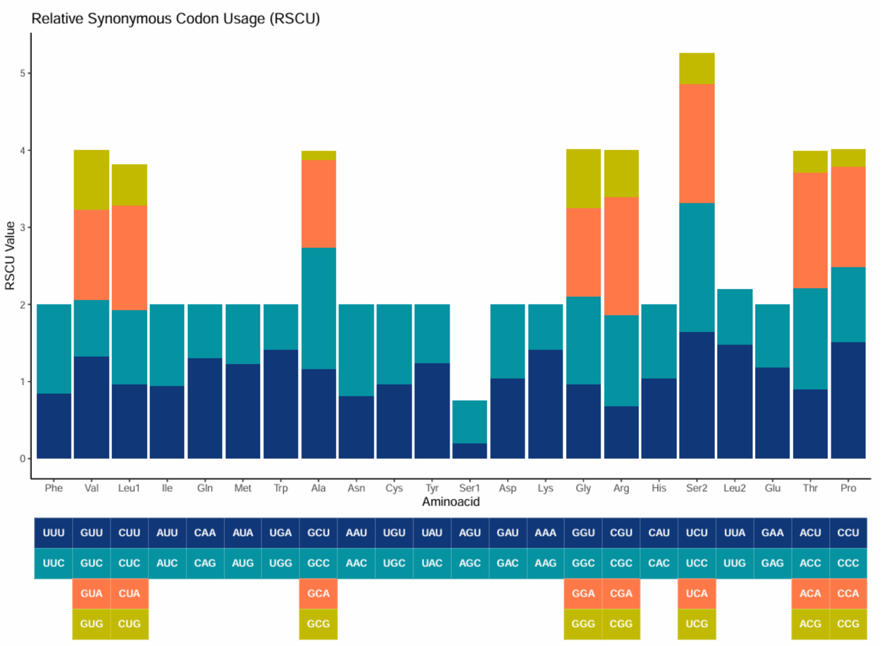

**Fig. C.1.** Relative Synonymous Codon Usage of P. pristis from Bonfim do Arari, Maranhão state, Brazil, western Atlantic Ocean.

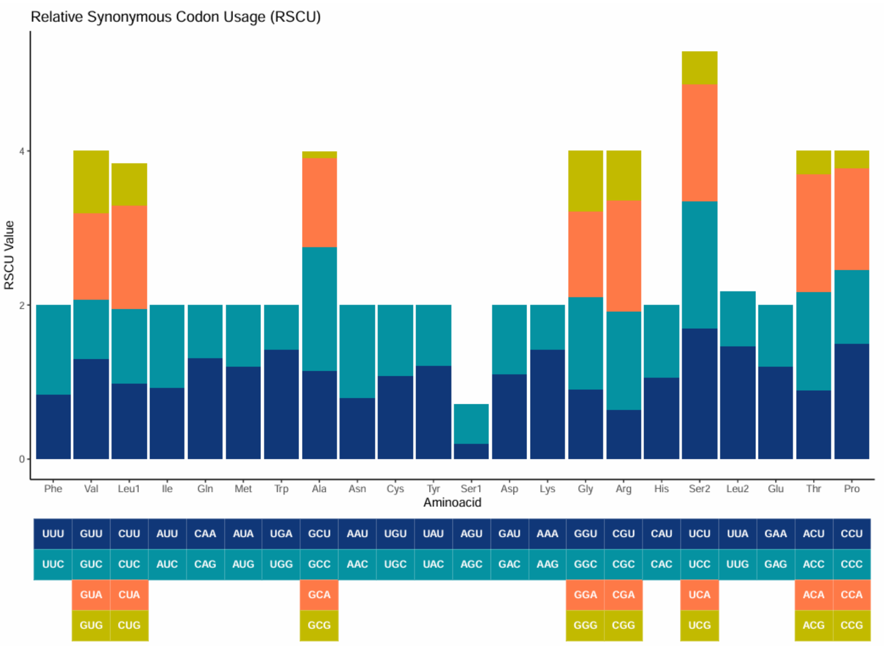

**Fig. C.2.** Relative Synonymous Codon Usage of *P. pristis* from Talara, Eastern Pacific in Peru.

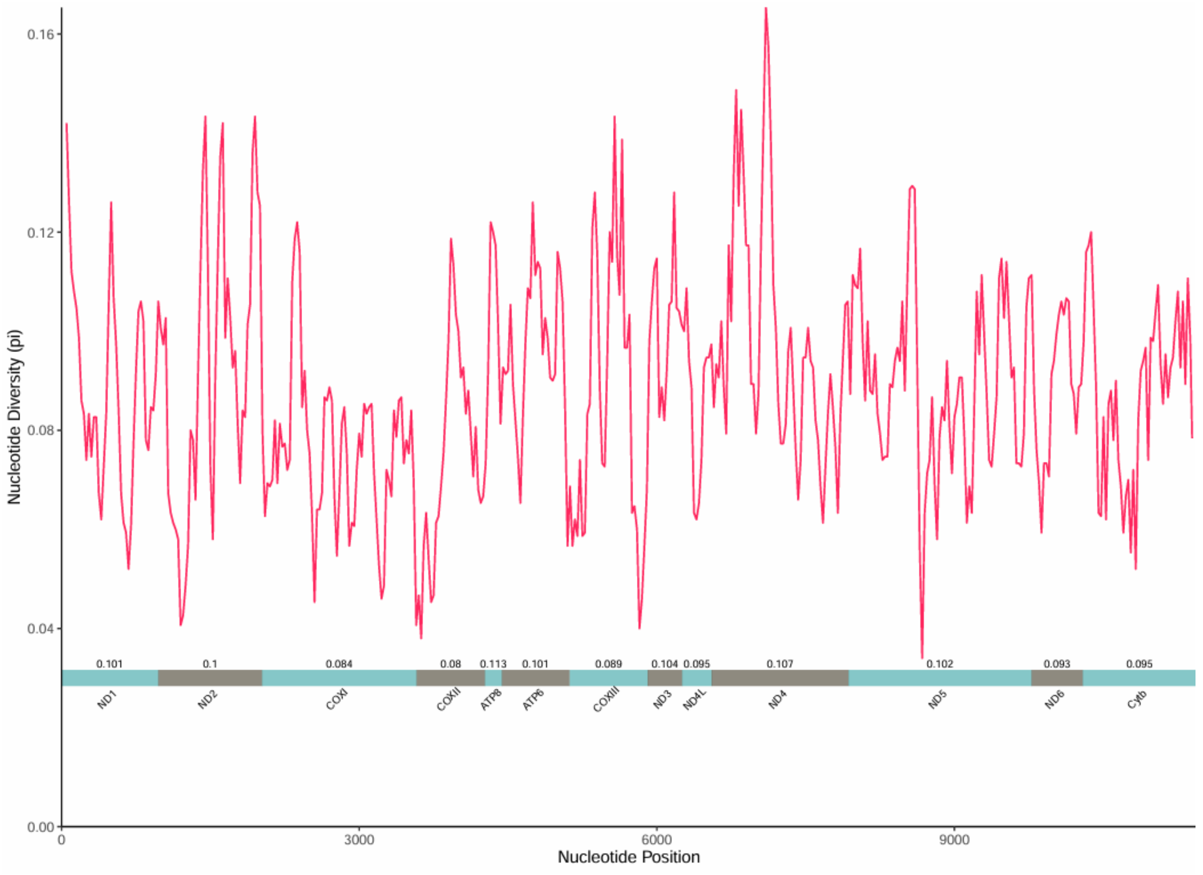

**Fig. C.3.** Sliding window-based nucleotide diversity plot for the genus *Pristis*. The red line represents the value of pi at each step of the analysis. The horizontal bars below represent the size of each PCG. The number above the bars shows the value of pi for 13 PCG individually.

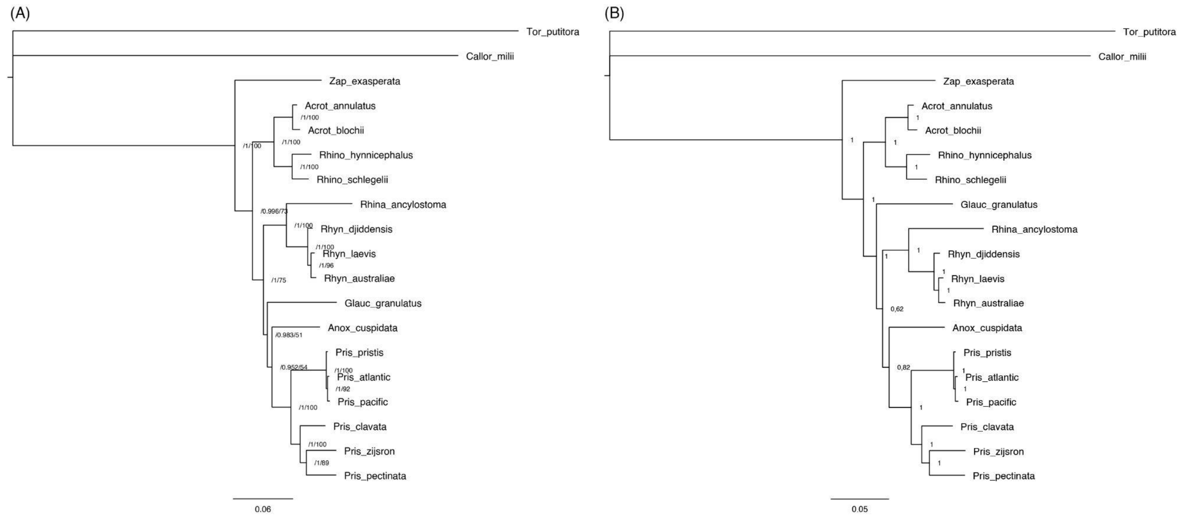

**Fig. C.4.** Phylogenetics tree for Rhinopristiforms showing the close relationship of samples of *Pristis pristis* from the Atlantic and Pacific coasts of South America to the existing sequence for a sample from Australia. (A) Support for nodes based on abayes approximation (left hand value) and Maximum Likelihood support generated using 1.000.000 pseudoreplicates in an UltraFast Bootstrap analysis (right hand value). (B) Bayesian Inference based on one million generations.

**Appendix D:** Primers design procedures and test with positive control

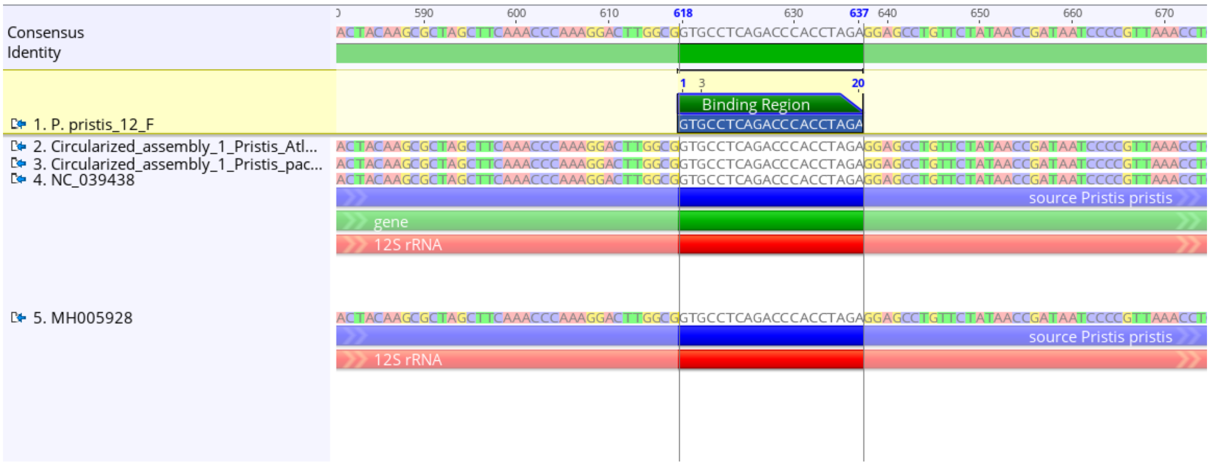

**Fig. D.1**. Alignment of the Forward primer sequence targeting a 12S fragment of *P. pristis* (Cooper et al. 2021) with two *P. pristis* mitogenomes available in GenBank, and the two mitogenomes provided in this study. No mismatches were found.

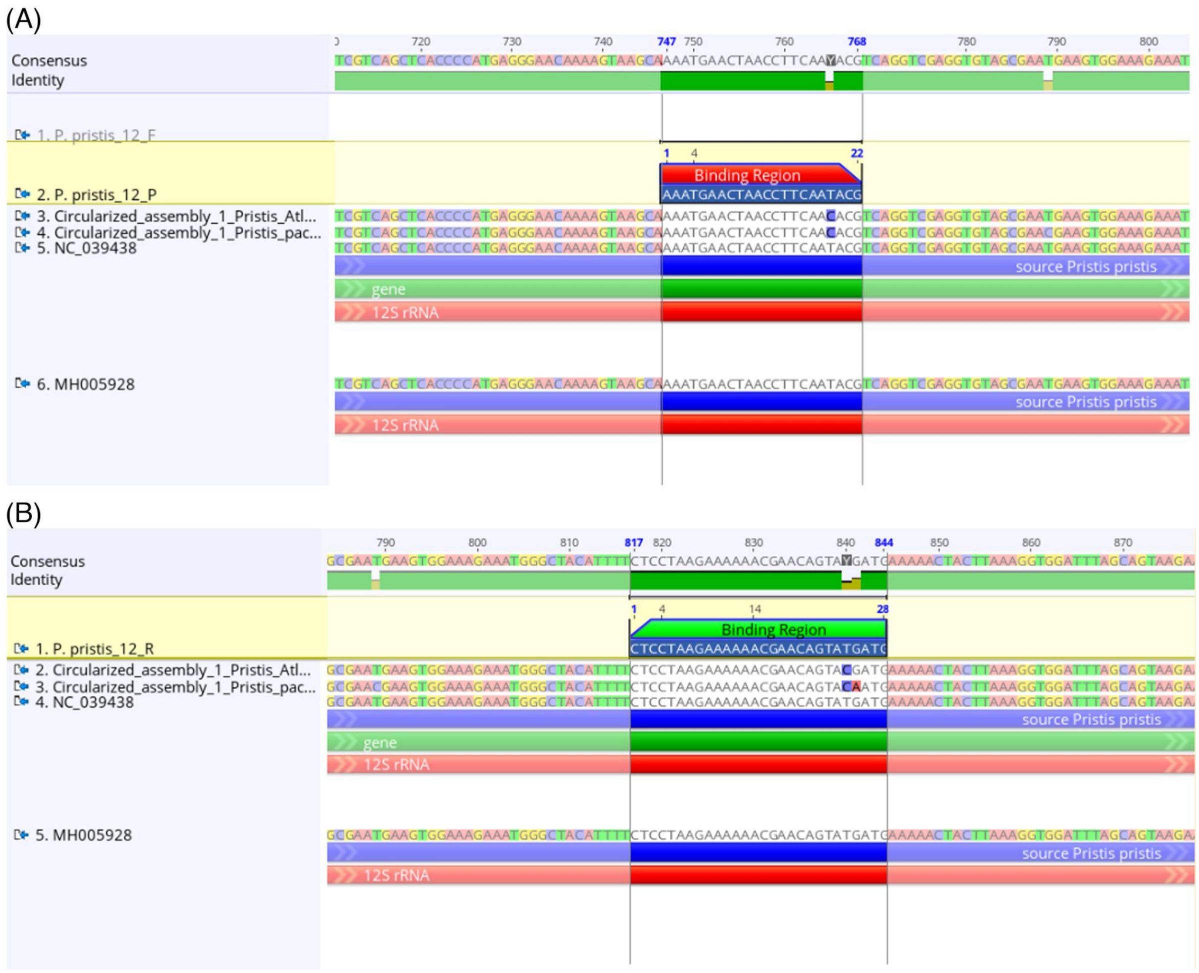

**Fig. D.2.** Local alignment for the existing 12S species-specific probe (A) and reverse primer (B) for *Pristis pristis* (Cooper et al. 2021) showing the mismatch of nucleotides for both the Western Atlantic and Eastern Pacific mitogenome lineages that prevent primer binding. Probe sequence (A) and reverse primer (B) were aligned with two Australian *P. pristis* mitogenomes available in GenBank, and the two new mitogenomes produced in this study.

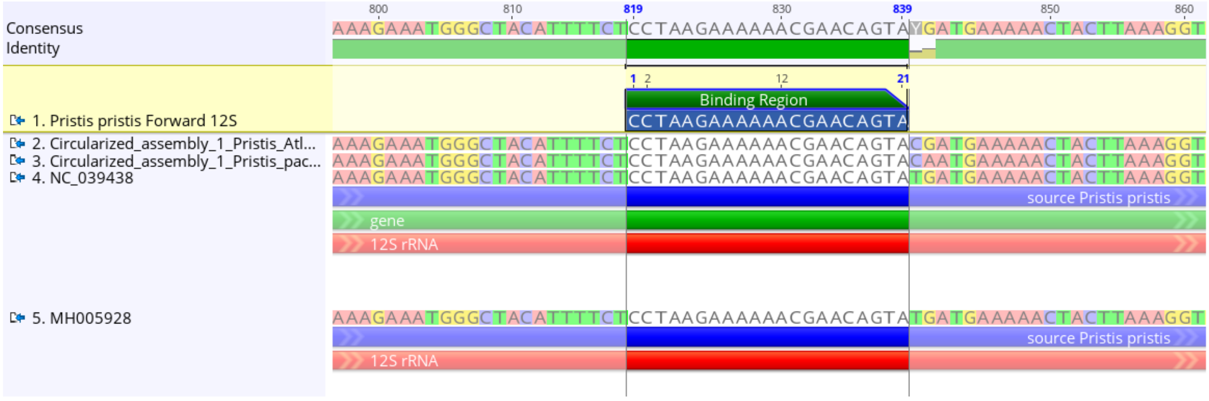

**Fig. D.3**. Alignment of the Forward primer sequence targeting a 12S fragment of *P. pristis* designed in this study with two *P. pristis* mitogenomes available in GenBank, and the two mitogenomes provided in this study. No mismatches were found.

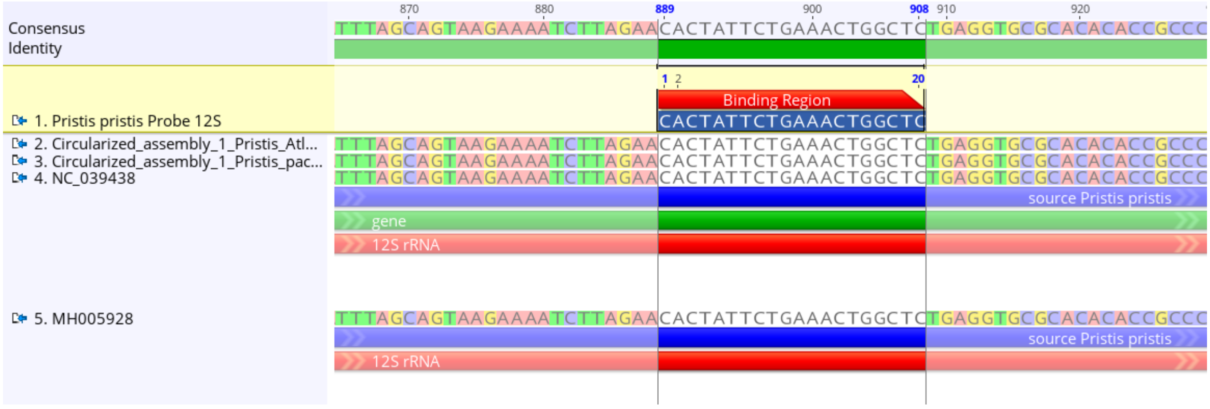

**Fig. D. 4**. Alignment of the Probe sequence targeting a 12S fragment of *P. pristis* designed in this study with two *P. pristis* mitogenomes available in GenBank, and the two mitogenomes provided in this study. No mismatches were found.

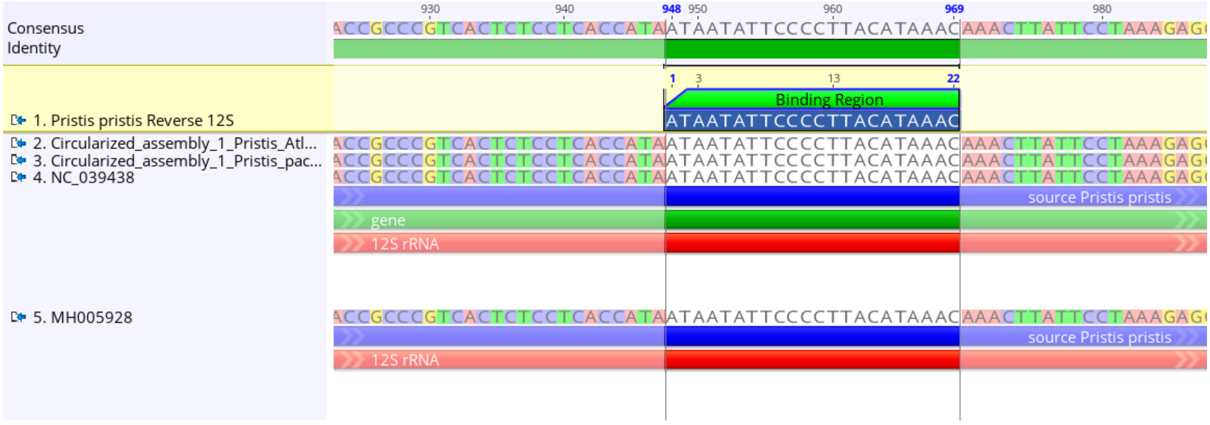

**Fig. D.5**. Alignment of the Reverse primer sequence targeting a 12S fragment of *P. pristis* designed in this study with two *P. pristis* mitogenomes available in GenBank, and the two mitogenomes provided in this study. No mismatches were found.

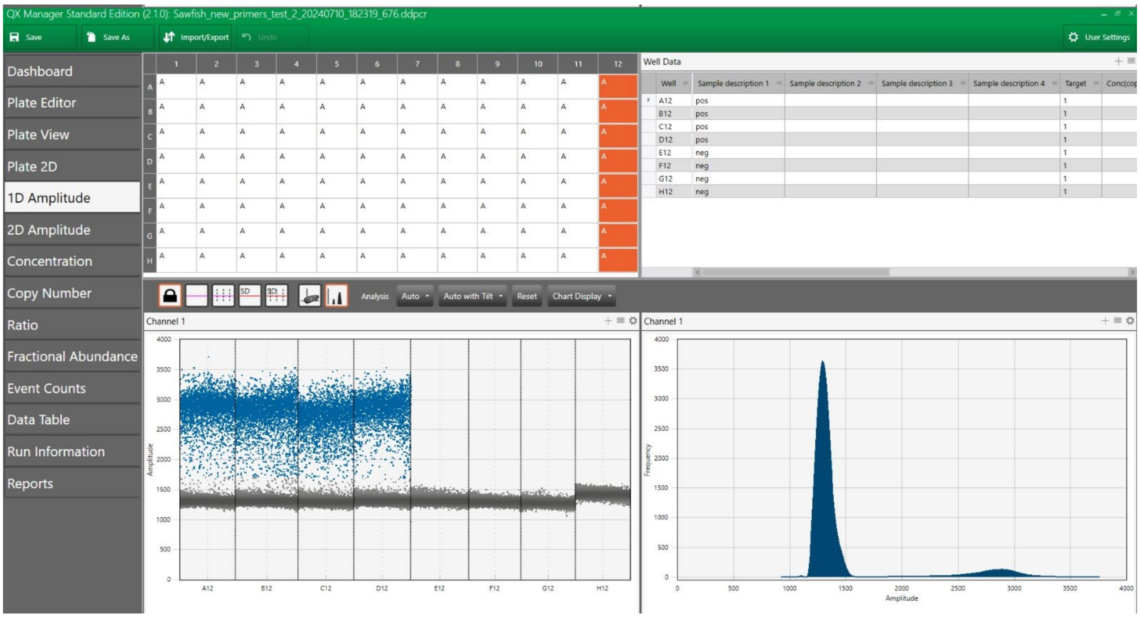

**Fig. D.6.** Comparison of success of ddPCR with positive and negative controls for the newly developed ddPCR 12S primers and probes for *P. pristis*. Positive controls: stomach samples replicate DNA extractions, 4 left hand lanes. Example negative controls: marine eDNA samples from northern Brazilian coastal waters, 4 right hand lanes.

Info about the analysed eDNA samples

ddPCR: 3 runs

1st run: I ran samples from the strip of plate 33 which had the DNA extracts from the sawfish and the strip 1 and 2 of plate 29. Only the wells C, D, E, F and G of plate 33 were positive (with very high levels) for both replicates. Everything else was negative. I didn't include specific negative controls, as were some on the strip from plate 33. All positive samples from plate 33 are coming from the same *Pristis pristis* stomach content.

2nd run: Still samples from plate 29. I ran samples from strip 3 to strip 8 with 2 replicates per samples, and then strip 8 with no replicate per samples. I included a strip with 4 positive controls made by diluted one of the positive samples from plate 33 (a 1:1000 dilution), and 4 negative controls. Only the 4 positives showed any DNA.

3rd run: Samples from plate 30. I ran samples from strip 1 to strip 9 and then strip 11 only with no replicate. I included a strip with 4 positive controls made by diluted one of the positive samples from plate 33 (a 1:1000 dilution), and 4 negative controls. Only the 4 positives showed any DNA.
